# CDK5 activity in retinal pigment epithelium contributes to gap junction dynamics during phagocytosis

**DOI:** 10.1101/2023.02.09.527850

**Authors:** Julia Fadjukov, Sophia Wienbar, Nemanja Milićević, Satu Hakanen, Maija Vihinen-Ranta, Teemu O. Ihalainen, Gregory W. Schwartz, Soile Nymark

## Abstract

Retinal pigment epithelium (RPE) at the back of the eye is a monolayer of cells with an extensive network of gap junctions that contributes to retinal health in a multitude of ways. One of those roles is the phagocytosis of photoreceptor outer segments. This renewal is under circadian regulation and peaks after light onset. Connexin 43 (Cx43) is the most predominantly expressed gap junction protein in RPE. In this study, we examine how gap junctions and specifically, Cx43 phosphorylation, contribute to phagocytosis in both human embryonic stem cell derived RPE and mouse RPE monolayers. We show that both Rac1 and CDK5 have differences in protein localization at different points in phagocytosis, and that by using their effectors, the capability of RPE for phagocytosis changes. CDK5 has not yet been reported in RPE tissue, and here we show that it likely regulates Cx43 localization and resulting electrical coupling. We find that gap junctions in RPE are temporally highly dynamic during phagocytosis and that regulation of gap junctions via phosphorylation is likely critical for maintaining eye health.

## Introduction

Gap junctions are intercellular channels that allow the direct passage of small ions and metabolites between adjacent cells. In vertebrates, these cell-cell junctions are composed of proteins known as connexins (Cx) that vary in their expression pattern between different tissues (Goodenough et al., 1996). Of the 21 connexin types found in the human genome, Cx43 is the most commonly expressed Cx isoform (Willecke et al., 2002). Cells of the retinal pigment epithelium (RPE) primarily express Cx43, but low levels of Cx36 have also been identified (Fadjukov et al., 2022; Malfait et al., 2001; Milićević et al., 2021).

To form a gap junction, six Cx proteins assemble into a hemi-channel, or connexon, that is then trafficked to the cellular membrane to dock with a hemichannel of the adjacent cell. Cx proteins are comprised of four hydrophobic transmembrane domains, two extracellular domains that help to dock the opposing cells, as well as three cytoplasmic domains: the amino (N)-terminal end, a loop connecting two of the transmembrane domains, and the carboxy (C)-terminal tail (Meşe et al., 2007; Pogoda et al., 2016; Willecke et al., 2002). Gap junctions are dynamic structures with reported half-lives of only 1-3 h (Falk et al., 2014; Laird et al., 1991; Lampe, 1994; Musil et al., 1990). Thus, regulation of the various stages of channel formation, trafficking, turnover, as well as gap junctional gating are likely to be crucial in the control of gap junction mediated cell-cell connectivity (Goodenough and Paul, 2009; Segretain and Falk, 2004). This intercellular communication can be regulated at multiple levels and timescales, including by changes in membrane voltage, pH, and post-translational modifications such as phosphorylation (Goodenough and Paul, 2009; Pogoda et al., 2016). The effect of phosphorylation on channel gating has been shown to be very specific, as phosphorylation on different residues by the same kinases can have opposite effects on the channel conductance (Lampe and Lau, 2004). Most of the known Cx phosphorylations target tyrosine and serine residues at the C-terminal end, and some of the implicated kinases for Cx43 include tyrosine kinase oncoprotein (Src), mitogen-activated protein kinase (MAPK), protein kinase A (PKA), protein kinase C (PKC), casein kinase 1 (CK1), pp60src kinase (for a review see (Lampe and Lau, 2004)) and cyclin dependent kinase 5 (Cdk5) (Qi et al., 2016). Thus far, Cdk5 has mainly been studied in the nervous system (Liu et al., 2008; Qi et al., 2016), but it has been shown to regulate various physiological and pathological functions in non-neuronal cells such as pancreatic cells and neutrophils (Contreras-Vallejos et al., 2012).

Clusters of the gap junction channels enable cell-cell communication that is believed to be vital for a wide range of physiological events, including synchronization of cellular membrane potentials, differentiation, proliferation, and tissue homeostasis support (Lampe, 1994; White and Paul, 1999). In addition, gap junctions have been implicated in regulation of phagocytosis in macrophages, although their role has been controversial (Anand et al., 2008; Dosch et al., 2019; Glass et al., 2013). This study addressed the potential role of gap junctions in a specialized phagocytosis process in RPE that aids the renewal of photoreceptors. This process is crucial for vision maintenance, and its molecular pathway is evolutionary conserved and under strict diurnal control (LaVail, 1976, 1980; Young, 1978). While photoreceptor outer segment (POS) phagocytosis is a continuous process, there is a bulk phagocytosis event occurring at light onset in a range of species (Bobu et al., 2006; Fisher et al., 1983; Flannery and Fisher, 1984; Ko et al., 2009; Nandrot et al., 2004; Strauss, 2005). The phagocytosis process involves coordinated tethering and internalization of the POS by the RPE with the help of surface receptors αvβ5 integrin and tyrosine kinase MerTK. The resulting phagocytic cup formation involves activation of RhoA family GTPases such as Rac1 and Cdc42, and rearrangement of cytoskeletal F-actin (Mao and Finnemann, 2012) and the ingested phagosomes are eventually digested through the endosomal-lysosomal pathway (Wavre-Shapton et al., 2014). Activation of various voltage-gated ion channels has also been implicated in the pathway, such as voltage-gated sodium (Na_V_) and calcium channels (Johansson et al., 2019; Korkka et al., 2019; Mamaeva et al., 2021; Müller et al., 2014).

In this study, we observe that Cx43 localizes close to opsin labeled POS particles during peak phagocytosis timepoints indicating that the localization of Cx43 connexons changes during the process. We investigate this relocalization further in cultured human embryonic stem cell derived RPE (hESC-RPE). Our results reveal that the change in gap junction localization is dependent on the Rac1 RhoA GTPase pathway. We demonstrate that the observed gap junction dynamics are regulated by phosphorylation of Cx43, particularly at the serine residue 279 (S279). Moreover, in this study we show that during phagocytosis, PKC regulates gap junctions and that their translocation can be prevented by inhibition of another kinase, Cdk5. The inhibition of Cdk5 results in higher junctional level of Cx43, leading to increased gap junctional coupling. This kinase has previously not, to our knowledge, been reported to be present in RPE tissue. Lastly, we use live imaging of mEGFP-tagged Cx43 in human induced pluripotent stem cell derived RPE (hiPSC-RPE) to observe gap junction localization in response to a Cdk5 inhibitor and POS challenge. Our data shows that inhibiting Cdk5 activity significantly promotes junctional localization of endogenous Cx43 and decreases the efficiency of POS phagocytosis.

## Methods

### Cell culturing

Human ESC lines Regea08/017 and Regea11/013 were cultured and spontaneously differentiated into RPE in floating cell clusters as previously described (Hongisto et al., 2017; Skottman, 2010; Vaajasaari et al., 2011). The mono-allelic endogenously mEGFP-tagged GJA1 WTC human induced pluripotent stem cell (iPSC) line AICS-0053-016 (Allen Cell Collection, Coriell Institute, Camden, NJ, USA) was also cultured and differentiated as above. Briefly, the pigmented areas were enzymatically dissociated using TrypLE Select (Invitrogen, UK) and seeded onto Collagen IV (5 µg/cm^2^, Sigma-Aldrich, St. Louis, MO) coated 24-well cell culture plates (Corning CelBIND; Corning, Inc., Corning, NY) at cell density 5.5 × 10^5^. The pigmented cells were then plated with a density of 2.5 × 10^5^ cells/cm^2^ onto collagen IV (C5533, Sigma-Aldrich, 10 µg/cm^2^) and Laminin 521 (LN521, Biolamina; 1.8 µg/cm^2^) coated hanging culture inserts (Millicell Hanging Cell Culture Insert, polyethylene terephthalate, 1.0 µm pore size, EMD Millipore, MA, USA) and maintained at 37°C in 5 % CO_2_. The inserts received Knock-Out Dulbecco’s modified Eagle’s medium (10829-018, Gibco) supplemented with 15 % Knock-Out serum replacement (10828-028, Gibco), 2 mM GlutaMax (35050-038, Gibco), 0.1 mM 2-mercaptoethanol (81350-010, Gibco), 1 % Minimum Essential Medium nonessential amino acids (1140050, Thermo Fisher Scientific), and 50 U/mL penicillin/streptomycin (from Cambrex BioScience, Walkersville,MD) and the medium was changed three times per week. Cells were allowed to mature for 8-12 weeks prior to experiments and their maturity was evaluated by immunostainings, transepithelial electrical resistance (TER) measurements and functional assays similarly to (Korkka et al., 2018).

The hESC lines of this study were obtained through a collaboration with Prof. Heli Skottman’s group at Tampere University, Finland. Tampere University, Faculty of Medicine and Health Technology, has the approval of the National Authority Fimea (Dnro FIMEA/2020/003758) to conduct research on human surplus embryos excluded from infertility treatments. The research is conducted under supportive statements of the Ethical Committee of the Pirkanmaa Hospital District to derive, culture, and differentiate hESC lines for research purposes (Skottman/R05116). New cell lines were not derived in this study.

### Sample preparation

To investigate the effect of Cdk inhibition, the AICS-mGJA1 and hESC inserts were incubated either with 100 µM roscovitine (Cat. no. 1332, R&D Systems) or equivalent amount of DMSO for 0 – 24h. In some of the experiments, the 24h incubation was followed by 0 – 90 min incubation with either 165 nM Phorbol 12-myristate 13-acetate (PMA, Cat. no. 1201, Tocris) or equivalent amount of DMSO. For monolayer patch clamp recordings and immunolabeling, the culture insert was removed from its holder and cut into smaller pieces. The cells were rinsed either with PBS (for immunolabeling) or with Ames’ solution (for patch clamp recordings). For live imaging of AICS-mGJA1, the incised insert was positioned in the Aireka coverslip cell chamber (Cat. no. SC15022, AirekaCells) by securing it under a slice anchor with the apical side facing the objective. The chamber was then filled with cell culture medium with or without blockers.

Mouse RPE from C57BL/6 mice was prepared for immunolabeling by euthanizing the animals by CO_2_ inhalation and cervical dislocation. The eyes were enucleated and bisected along the equator, after which the eyecups were sectioned in Ames’ solution buffered with 10 mM HEPES and supplemented with 10 mM NaCl, pH was adjusted to 7.4 with NaOH (Sigma-Aldrich). The retina was gently removed from the eyecup leaving the RPE firmly attached to the eyecup preparation. All mice were between 4 – 10 weeks old and on a mixed C57BL/6 background. Both male and female mice were used in this study. The mice were reared in a 12 h light/dark cycle. All procedures were approved by the Animal Care and Use Committee at Northwestern University and in accordance with the ARVO Statement for the Use of Animals in Ophthalmic and Vision Research and Finland Animal Welfare Act 1986.

### RNA isolation, cDNA synthesis and mRNA quantification

Total RNA was isolated from hESC RPE using the RNeasy mini kit (Qiagen, Valencia, CA, USA) according to the manufacturer’s instructions in which we performed the optional DNAse I (Qiagen, Valencia, CA, USA) incubation step. The elutes had RNA yield between 41-61.6 ng / μl / insert. Complementary DNA was synthesized from 200 ng of total RNA using oligo(dT)_12-18_ primed reactions with Superscript III reverse transcriptase (Life technologies, Waltham, MA, USA). The synthesized cDNA was amplified with transcript-specific, intron-spanning primers with PCR amplification cycles optimized for product quantification. Primer sequences and PCR conditions are provided in the Supplementary figure 1. PCR products were electrophorized on 2% agarose gels containing GelRed® (Biotium, Fremont, CA, USA) and images were captured using the ChemiDoc(tm) MP System (Bio-Rad laboratories, Orlando, FL, USA). Bands were quantified using Image Lab (version 5.2, Bio-Rad Laboratories, Orlando, FL, USA). Band intensities for each gene were divided with the expression value of the reference gene *GAPDH*. The normality of the data was assessed with Shapiro–Wilk normality test. The mRNA levels were analyzed statistically with 2-tailed unpaired Student’s t-test. The alpha level was 0.05 for all statistical tests mentioned in the methods.

### Phagocytosis assay for cultured and mouse RPE

The porcine POS particles were isolated and purified as previously described (Johansson et al., 2019; Mao and Finnemann, 2013). Briefly, the eyecups obtained from a slaughterhouse were opened using a scalpel and retinas were removed with forceps under dim red light. The collected retinas were agitated in 0.73 M sucrose phosphate buffer, filtered and separated in sucrose gradient with ultracentrifugation (Optima ultracentrifuge, Beckman Coulter, Inc., Brea, CA) at 112,400 x g for 48 min at +4 °C. The collected POS layer was centrifuged 3000×g for 10 min at +4 °C and stored in 73 mM sucrose phosphate buffer at -80 °C prior to experiments.

The purified POS particles were fed to the hESC-derived RPE cells in a KO-DMEM medium supplemented with 10% fetal bovine serum (FBS) and incubated for either 15 min, 30 min or 2 h at +37 °C in 5% CO_2_. Phagocytosis was studied live by placing the AICS-mGJA1 RPE inserts in the Aireka imaging chamber after a 20 min POS challenge as described in the sample preparation chapter. In the pharmacological experiments, the samples were preincubated for either 3 h with Rho/Rac/Cdc42 Activator I (1 µg/ml, Catalog number CN04, Cytoskeleton Inc), 30 min with MEK inhibitor PD 98059 (50 µM, Cat. no. 513001, Sigma-Aldrich), 30 min with the PKC activator Phorbol 12-myristate 13-acetate (PMA) (165 nmol) or 24 h with the CDK5 inhibitor Roscovitine (100 µM) or for the longest equivalent time with DMSO (Control). The compounds and DMSO were present for the duration of the phagocytosis experiment unless otherwise indicated in the figures. Then the monolayers were washed with PBS and fixed with PFA according to the immunostaining protocol. Phagocytosis was studied in vivo by preparing the mouse eyes under dim red light either at light onset or 1.5 h, 2 h or 6 — 10 h after it.

### Quantification of POS particles in hESC-derived RPE

To detect and quantify POS particles, random fields chosen blindly by ZO-1 labeling were imaged from 3 different samples in each condition with Nikon A1R. The images were processed with a Gaussian function and the obtained maximum intensity z-projection was binarized using a global threshold. The number of POS particles was then analyzed from the images converted to mask using the particle analysis of ImageJ (Schneider et al., 2012). Analysis of internalized POS particles was carried out from maximum intensity projections of xy-slices as has been previously described (Viheriälä et al., 2021).

Each phagocytosis experiment was repeated at least three times and the images were pooled together. The normality of the data was tested by using Shapiro–Wilk normality test. To compare differences between roscovitine treated and control samples, the significance was analyzed using the Mann Whitney U test. To compare differences between Rac/Rho/Cdc42 Activator I treated and control samples, the significance was analyzed using the 2-tailed unpaired Student’s t-test.

### Immunolabeling

All samples were fixed for 15 min with 4% paraformaldehyde (pH 7.4;15713 Electron Microscopy Sciences). After three 5 min washes with phosphate buffered saline (PBS), samples were permeabilized for 15 min in 0.1% Triton X-100 in PBS (Sigma-Aldrich) and blocked with 3% bovine serum albumin/PBS (BSA; Sigma-Aldrich) for 1 h at RT.

The following primary antibodies were used in this study: Connexin 43 (Cx43) 1:200 (C6219, Sigma-Aldrich, RRID:AB_476857), Opsin 1:200 (O4886, Sigma-Aldrich, RRID:AB_260838), Na_V_1.4 1:200 (ASC-020, Alomone labs, RRID:AB_2040009), ZO-1 1:50 (33-9100, Thermo Fisher Scientific, RRID:AB_2533147), Rac1 1:100 (610650, BD Biosciences RRID:AB_397977), Phospho-Connexin43 Ser373 (S373) 1:200 (PA5-64670, Thermo Fisher Scientific, RRID:AB_2662693), Phospho-Connexin43 Ser279 (S279) 1:200 (PA564640, Thermo Fisher Scientific, RRID:AB_2662450). All primary antibodies were diluted in the blocking solution and incubated on the tissue for 1 h at RT. After three PBS washes and 1 h RT incubation with the secondary antibodies that were all diluted 1:200 in 3% BSA in PBS: goat anti-rabbit Alexa Fluor 568 (A-11011) and donkey anti-mouse Alexa Fluor 488 (A10037) (Thermo Fisher Scientific), the actin cytoskeleton was labeled with a direct phalloidin ATTO 643 (Cat no. AD643-81, ATTO-TECH) 1:100 or ATTO 488 conjugate (Cat no. 49409, Sigma-Aldrich) 1:100. All samples were mounted with Prolong Diamond (P36961, Thermo Fisher Scientific).

### Immunogold labeling and electron microscopy

The hESC RPE monolayers were fixed and prepared for immunogold labeling as previously described (Johansson et al., 2019: Fadjukov et al., 2022). After washing three times with PBS, samples were fixed with periodate-lysine-paraformaldehyde (PLP) for 2 hours at room temperature. Tissue was then treated with 0.01% saponin and 0.1% BSA in 0.1 M phosphate buffer (PB) (pH 7.4, Buffer A). The primary antibody concentrations were quadrupled compared to immunolabeling, diluted in Buffer A and incubated for 1h. The secondary antibody (1.4 nm nanogold-conjugated polyclonal Fab’ fragment of goat anti-rabbit IgG; Nanoprobes.com, Yaphank, NY) was diluted to 1:50 in Buffer A and was incubated for 1 hr. Following washes with Buffer A and phosphate buffer (PB), tissue was post-fixed for 10 min at room temperature with 1% glutaraldehyde in PB. Then, it was quenched for 5 min in 50 mM NH_4_CL in PB, followed by washes in PB. Samples were treated with HQ-silver (Nanoprobes.com) for 5 min in dark and washed with water. Then, they were gold toned with 2% sodium acetate 3 × 5 min at RT, 0.05% gold chloride 10 min at +4 °C, 0.3% sodium thiosulphate 2 × 10 min at +4 °C, washes with water. Next, the samples were reduced for 1 h at +4 °C with 1% osmium tetroxide in 0.1 M phosphate buffer, dehydrated with graded series of ethanol (70%, 96%, 100%), and stained with 2% uranyl acetate. Finally, the labeled samples were embedded in Epon (TAAB Embedding resin, medium, TAAB Laboratories Equipment Ltd, Berks, UK) and polymerized. For sectioning, samples were sliced at 200 nm intervals with a microtome (Leica ultracut UCT ultramicrotome, Leica Mikrosysteme GmbH, Austria), ensuring they were oriented perpendicular to the monolayer. Slices were placed on carbon-coated single-slot grids and imaged at 80 kV voltage with JEOL JEM-1400 transmission electron microscope (JEOL Ltd., Tokyo, Japan) equipped with bottom-mounted Quemesa CCD camera (4008 × 2664 pixels).

Mouse RPE was dissected from mixed C57BL/6 mice and fixed in 4% paraformaldehyde and 2.5% sucrose in 0.1 M PB overnight. Coronal sections of the eye were immersed in 2.3 M sucrose in PB and rotated at +4°C for 4h. Then, tissue was frozen in liquid nitrogen and thin cryosections were cut with a Leica EM UC7 cryoultramicrotome (Leica Microsystems, Vienna, Austria). The sections were picked on Butvar – coated nickel grids. Grids were then incubated in 2% gelatine in PB for 20 min then in 0.1% glycin-PB for 10 min. Blocking with Protein A/G Gold conjugates (Aurion, The Netherlands) for 15 min. Washing and antibody incubation steps were performed in 0.1% BSAc (Aurion, The Netherlands) in PBS. Primary antibody against Cx43 (1:50) was incubated for 45 min, controls used only PBS. Then incubation with protein A conjugated 10 nm gold (Cell Microscopy Core, University Medical Center Utrecht, The Netherlands) occurred for 30 min. Finally, grids were stained with neutral uranyl acetate (UA) and coated with 2% methyl cellulose containing 0.4 % UA. Sections were examined with a Tecnai G2 Spirit 120 kV transmission electron microscope (FEI, Eindhoven, The Netherlands) and images were captured by a Quemesa CCD camera (Olympus Soft Imaging Solutions GMBH, Münster, Germany) using RADIUS software (EMSIS GmbH, Münster, Germany).

### Confocal microscopy and image processing

Confocal microscopy of the RPE was carried out using Nikon A1R laser scanning confocal microscope mounted in inverted Nikon Ti–E (Nikon Instruments Europe BV, Amsterdam, Netherlands) with a Plan-Apochromat 60x/1.4 oil immersion objective by acquiring 1024×1024 sized pixel 3D z-stacks. The following excitation laser lines and emission filters were used: 488 nm and 525/50; 561 nm and BP595/50 nm; and 633 nm and 700/75 nm. The laser intensities and detector sensitivities were adjusted for each sample to optimize the image brightness and to avoid saturation. For live imaging experiments, the mEGFP-tagged mGJA1 AICS monolayers in Aireka imaging chambers were kept at 37°C and a Z-stack was obtained every 30 min with Zeiss780 Zeiss LSM780 LSCM on inverted Zeiss Cell Observer microscope (Zeiss, Jena, Germany) by using Zeiss C Apo 63x/1.20 WD objective and 488 nm laser line for a total duration of 90 – 150 min.

The Cx43 foci analysis was performed by ImageJ particle analysis. The differences between the groups were analyzed by Kruskal-Wallis test and the subsequent post-hoc pairwise comparisons performed with the IBM SPSS Statistics for Windows, version 26 (IBM Corp., Armonk, N.Y., USA. Final figure panels were assembled using Adobe Illustrator (Adobe Systems, San Jose, USA).

### Patch clamp recordings

Currents were recorded from mature hESC-derived RPE monolayers with the standard patch-clamp technique in whole-cell configuration. Patch pipettes (5–7 MΩ, BF120-69-10, Sutter Instruments) were filled with an intracellular solution composed of (in mM): 125 K-aspartate, 10 KCl, 1 MgCl2, 10 HEPES, 1 CaCl2, 2 EGTA, 4 Mg-ATP and 0.5 Tris-GTP (277 mOsm; pH ∼7.15 with KOH). During all recordings, the cells were perfused with Ames’ solution (Sigma-Aldrich) buffered with 10 mM HEPES and supplemented with 10 mM NaCl and 5 mM TEA-Cl. The pH was adjusted to 7.4 with NaOH and the osmolarity set to ∼ 295 mOsm. All recordings were made in current-clamp mode with a 2-channel patch-clamp amplifier (MultiClamp 700B, Molecular Devices). To analyze the input resistance, we applied a series of injected current pulses from -100 to 100 pA for hESC-RPE and measured the changes in the membrane potential. Series resistance was not compensated. In paired recordings, two adjacent neighbor cells were patched simultaneously from monolayers incubated either with DMSO (control) or 100 µM roscovitine for 24 h at 37°C. When input resistance was analyzed from the POS particle treated hESC-RPE, the inserts were kept in the incubator with POS for either 30 min or 2 h prior to placing the insert in the recording chamber.

Dissections of mouse RPE monolayers for patch clamp recording were performed under IR light in carbogenated AMES (A-1372-25; US Biological Life Sciences) media buffered with sodium bicarbonate as previously described (Fadjukov et al., 2022). Membrane voltage was recorded in whole-cell patch clamp configuration using current clamp mode (MultiClamp 700B; Molecular Devices). Patch pipettes were 5-7 MΩ and filled with a potassium based internal containing (in mM) 125 K-aspartate, 10 KCl, 1 MgCl2, 10 HEPES, 1 CaCl2, 2 EGTA, 4 Mg-ATP, and 0.5 TrisGTP (277 mOsm; pH ∼7.15 with KOH). Current injections were performed between -500 and +1000 pA and membrane potential and cell input resistance was measured. Series resistance was not compensated.

### Patch clamp data analysis

The offline analysis was performed with Clampfit software (Molecular Devices) or with a custom open-source MATLAB package (GitHub - SchwartzNU/SymphonyAnalysis). The coupling coefficient was calculated as change in voltage in the recorded cell over change in voltage in the stimulated cell. The normality of the data was tested by using Shapiro–Wilk normality test and the effect of the roscovitine was analyzed with 2-tailed unpaired Student’s t-test. All statistical tests were performed with the IBM SPSS Statistics for Windows, version 26 (IBM Corp., Armonk, N.Y., USA) and the final figures were assembled in Adobe Illustrator.

## Code Availability

All code used in analysis will be made available upon request.

## Data Availability

All data will be made available upon request.

## Results

### Gap junctions translocate during POS phagocytosis

In this study, we explored the role of gap junctions in POS phagocytosis. First, we examined the localization of Cx43 and opsin proteins in mouse RPE wholemounts during peak phagocytosis (light onset and up to 2 h post) and control timepoints (light onset + 10 h) (Figure 1a). It is worth noting here that phagocytosis *in vivo* is a continuous process, with a burst occurring at light onset in a variety of species (Ko et al., 2009; Nandrot et al., 2004; Strauss, 2005). Our immunolabeling results demonstrated that at light onset (Figure 1b) or 2 h post (Figure 1c) Cx43 localized together with the bound POS particles, as shown in the maximum intensity z-projections of the apical confocal sections (Figure 1b,c;). Outside peak phagocytosis time points, Cx43 was primarily localized at specific foci at the junctions (Figure 1d). In addition, the number of Cx43 positive foci was found to increase in the RPE during phagocytosis (n of fields 9) compared to control timepoints, indicating reorganization of connexons (n of fields 11) (p-value = 0.0097) (Figure 1e). Thus, during peak phagocytosis Cx43 relocalized from the cell periphery to the center of the cell and partially colocalized with POS particles.

**Figure 1:**
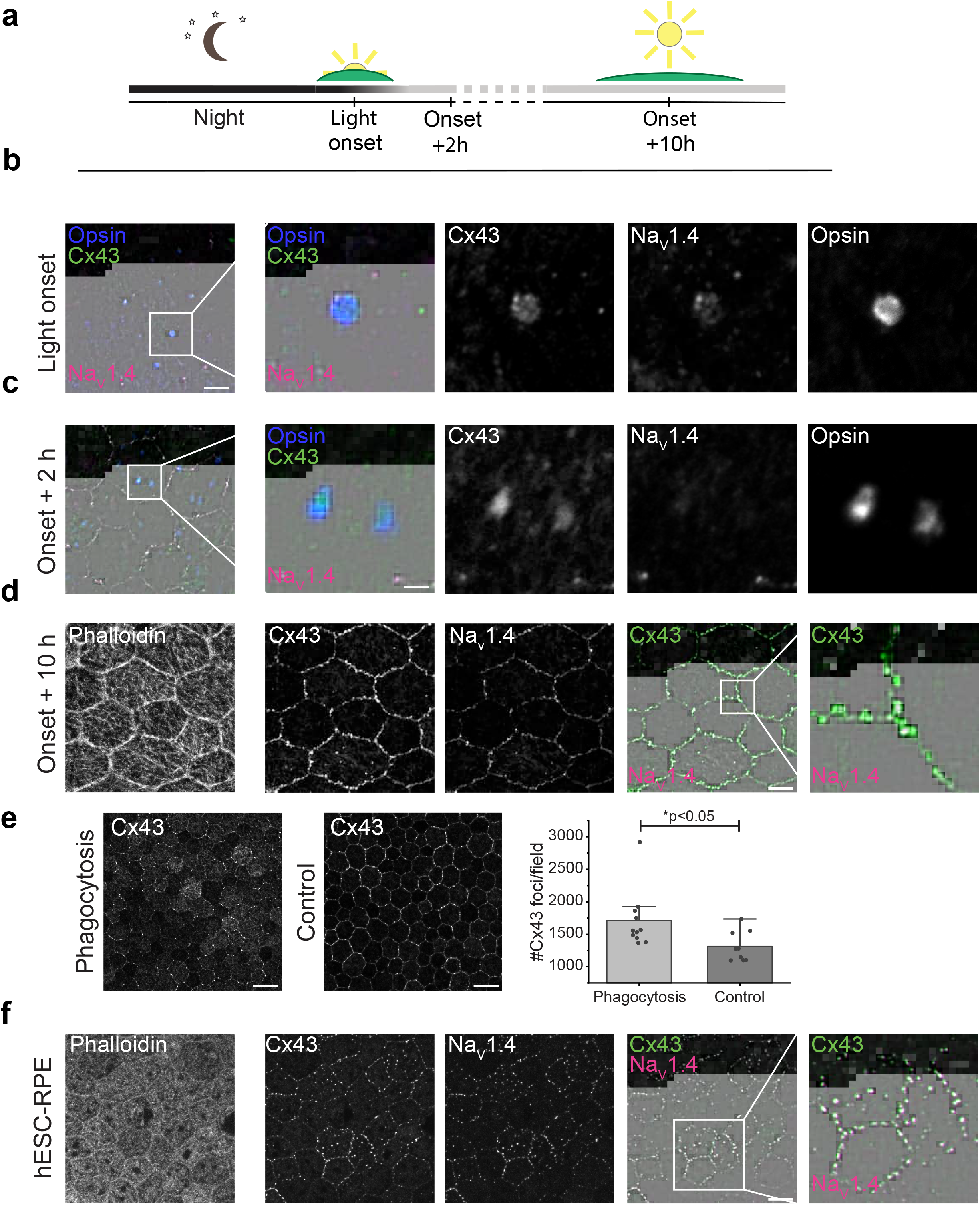
Cx43 and NaV1.4 immunolabeling in the presence and absence of POS phagocytosis. **a**. Schematic of the mouse eye dissection time points relative to the diurnal cycle. **b**. Mouse RPE dissected at light onset, and stained for Cx43 (*green*), NaV1.4 (*magenta*) and POS (opsin, *blue*). The maximum intensity z-projection includes sections from the apical side of RPE. Scalebar 10 µm. **c**. Same as **b**, however the RPE was dissected 2 hours post light onset. Scalebar 10 µm. **d**. RPE was dissected 10 hours post light onset, and stained for phalloidin (left), Cx43 (*green*), and NaV1.4 (*magenta*). The maximum intensity z-projection includes sections through the whole RPE. Scalebar 10 µm. **e**. Quantification of Cx43 positive foci in mouse RPE during phagocytosis (n = 9 fields) and control timepoints (n =11 fields). Data are shown as mean±SD. **f**. hESC RPE stained for Cx43 (*green*), NaV1.4 (*magenta*), and phalloidin (left). Scalebar 10 µm.

We had previously found that Na_V_ channels are involved in phagocytosis and undergo translocation during the process (Johansson et al., 2019). The labeling pattern of Cx43 was reminiscent of the labeling of a specific subtype of voltage-gated sodium channels Na_V_1.4. Moreover, Cx43 and Na_V_ channels have been shown to interact in cardiac myocytes (for review see (Veeraraghavan et al., 2014). Therefore, we investigated this plausible interaction in RPE by co-immunolabeling Cx43 and Na_V_1.4 during phagocytosis and in control conditions. Interestingly, the results showed a strong co-localization, and this phenomenon was observed in both hESC-derived and mouse RPE (Figure 1b,c,f). During phagocytosis, Na_V_1.4 was also found to localize with opsin, which was most evident at light onset. This result indicates that Cx43 and Na_V_1.4 might translocate together, suggesting that they interact in the phagocytosis process.

To investigate the time course of gap junction dynamics in more detail *in vitro*, we performed immunolabeling at several time points from 0 to 2 h during POS phagocytosis in hESC-RPE. We labeled Cx43 without the presence of POS (n = 20 fields) or at 15 min (n = 22 fields), 30 min (n = 21 fields), or 2 h (n = 23 fields) after the POS challenge (Figure 2a). Again, Cx43 was found to mainly localize into the junctions in control conditions, but its localization dramatically changed during the phagocytosis process. After 30 min, the labeling appeared more in the center of the cell in the maximum intensity z-projection as opposed to in the junctions; and at 2 h, the labeling was much more diffuse overall. This pattern was observed across three separate experiments, where the number of Cx43 positive foci was quantified for each timepoint (Figure 2a, right). The Kruskal-Wallis test revealed statistically significant Cx43 foci number changes during phagocytosis (p-value = 4.79 × 10^−11^). The subsequent post-hoc pairwise comparisons showed no statistical difference between no POS and 15 min (p-value = 0.991), but a significant increase in Cx43 labeling between no POS and 30 min groups (p-value = 0.033) as well as a significant decrease between no POS and 2h phagocytosis samples (p-value = 0,000004). Thus, it appeared that initial POS binding did not change Cx43 localization, but that as phagocytosis progressed, Cx43 was translocated from the cell-cell junctions.

**Figure 2:**
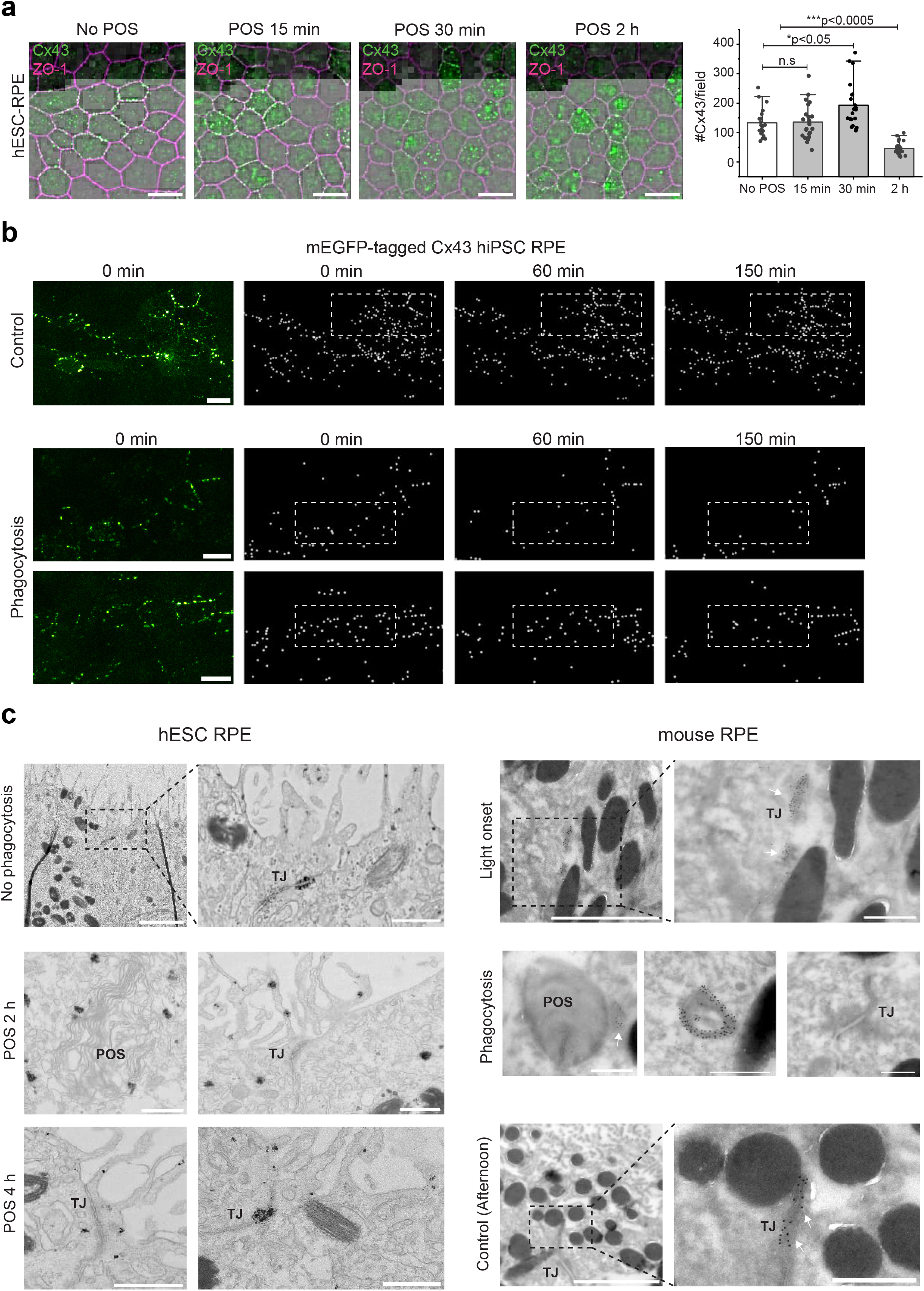
Translocation of Cx43 coincides with phagocytosis. **a**. Cx43 (*green*) and ZO-1 (*magenta*, tight junction marker) staining at different time points post application of POS particles. Quantification of Cx43 fluorescent foci per field at each time point (n = 3 experiments) is shown to the right. Data are shown as mean±SD. Scalebars 15 µm. **b**. Maximum intensity z-projections of Z-stacks from live imaging of hiPSC-RPE that expressed mEGFP-tagged Cx43 proteins without POS (control, top) or starting at 20 min after incubation with POS (phagocytosis). The Cx43 foci are shown in the time series as particles after local maxima detection, with example areas highlighted by dashed boxes. Scalebars 10 µm. **c**. EM images from hESC (left) and mouse (right) RPE at various time points in the phagocytic process labeled with immunogold for Cx43 and highlighted by white arrows in mouse RPE. Marked in the images are the tight junctions (TJ) and POS particles (POS). An internalized annular gap junction is shown in the middle of the mouse phagocytosis image. Scalebars 2 µm in lower magnification images (with dashed boxes) and 500 nm in others.

In order to examine Cx43 dynamics during the phagocytic process, we generated a hiPSC-RPE cell line that endogenously expresses mEGFP-tagged Cx43 and performed live cell imaging (Figure 2b). Prior to imaging, cells were incubated for 20 min with medium (control) or medium containing purified porcine POS particles (phagocytosis). The acquired confocal z-stack time-lapse videos showed that gap junctions are surprisingly dynamic in RPE (Supplementary video 1, 2, 3) but that in control conditions, EGFP-Cx43 positive foci were found primarily in the cell periphery. In contrast, after the POS challenge, a portion of the Cx43-positive puncta was found to disappear suggesting internalization and disassembly of EGFP-Cx43 tagged connexons. This correlates with the data collected from fixed cells, which showed reduction in the number of Cx43 positive foci at the 2 h mark (Figure 2a).

Next, we wanted to investigate the possible direct interplay and colocalization of Cx43 and POS particles in higher detail. This was achieved by conducting immuno-EM from hESC- and mouse RPE (Figure 2c). We wish to highlight that for cultured RPE, the samples could be processed by pre-embedding immunogold labeling yielding stronger and more robust signals. However, for mouse RPE we needed to use a technique that better preserves the tissue ultrastructure, albeit at the expense of signal strength and robustness, and thus opted for labeling cryosections. (Jones, 2016). Without phagocytosis (hESC) or at light onset (mouse), we found that gold nanoparticle labeled Cx43 were present at the cell-cell junctions. However, 2 h after phagocytosis challenge (hESC) or 1.5 h after light onset (mouse), the labeling was no longer consistently identified at the junctional area. Moreover, Cx43 was found to localize around ingested phagosomes or in structures that resembled internalized double-membrane structures known as annular gap junctions (Jordan et al., 2001) (Figure 2c). After 4 h (hESC) or afternoon control timepoints (mouse), the Cx43 labeling had partially or fully returned to the cell-cell junctions. Thus, in both hESC and mouse, the timing of Cx43 translocation correlated with the time points of phagocytosis. When we analyzed input resistance values after POS challenge in hESC-RPE (p-value = 0.588, Kruskal Wallis Test) or at various points of the diurnal cycle in mouse RPE, the overall input resistance levels were not found to remarkably change, yet we occasionally found cells with much higher input resistance values (Supplementary Figure 2). Cells with reduced gap junction connectivity would be expected to have a higher input resistance, thus these few examples might have captured the period where Cx43 connexons were internalized.

### Cx43 translocation is regulated by phosphorylation at serine 279

Phosphorylation is a known regulator of gap junctions, therefore, we investigated the possibility that phosphorylation of Cx43 occurs during phagocytosis. We investigated two potential phosphorylation sites, serine 373 (S373) and serine 279 (S279) (Figure 3a) because previous literature demonstrated that these sites were phosphorylated along similar time scales in response to injury or growth factors that we observed with phagocytosis (Lastwika et al., 2019; Solan and Lampe, 2014, 2018). Immunolabeling for Cx43 phosphorylated at S373, showed that this phosphorylated form was at some, but not most junctions in hESC-RPE. In addition, we observed no difference in the levels of phosphorylation with the addition of POS and the induction of phagocytosis (Figure 4a). However, the labeling pattern for a phosphorylated S279 specific Cx43 antibody was more dynamic: this serine modification was not detected, or detected at minimal levels, in control conditions, but the labeling increased during phagocytosis (Figure 4a). The signal was observed starting at 15 min after POS phagocytosis and its level was strongest after 30 min of POS incubation. The S279 labeling was no longer detected in the cell-cell junctions after 2 h of POS incubation (Figure 3a). Interestingly, during phagocytosis, the labeling was present at both the junctions and in the apical space, indicating that perhaps both gap junctions and hemichannels were being phosphorylated.

**Figure 3:**
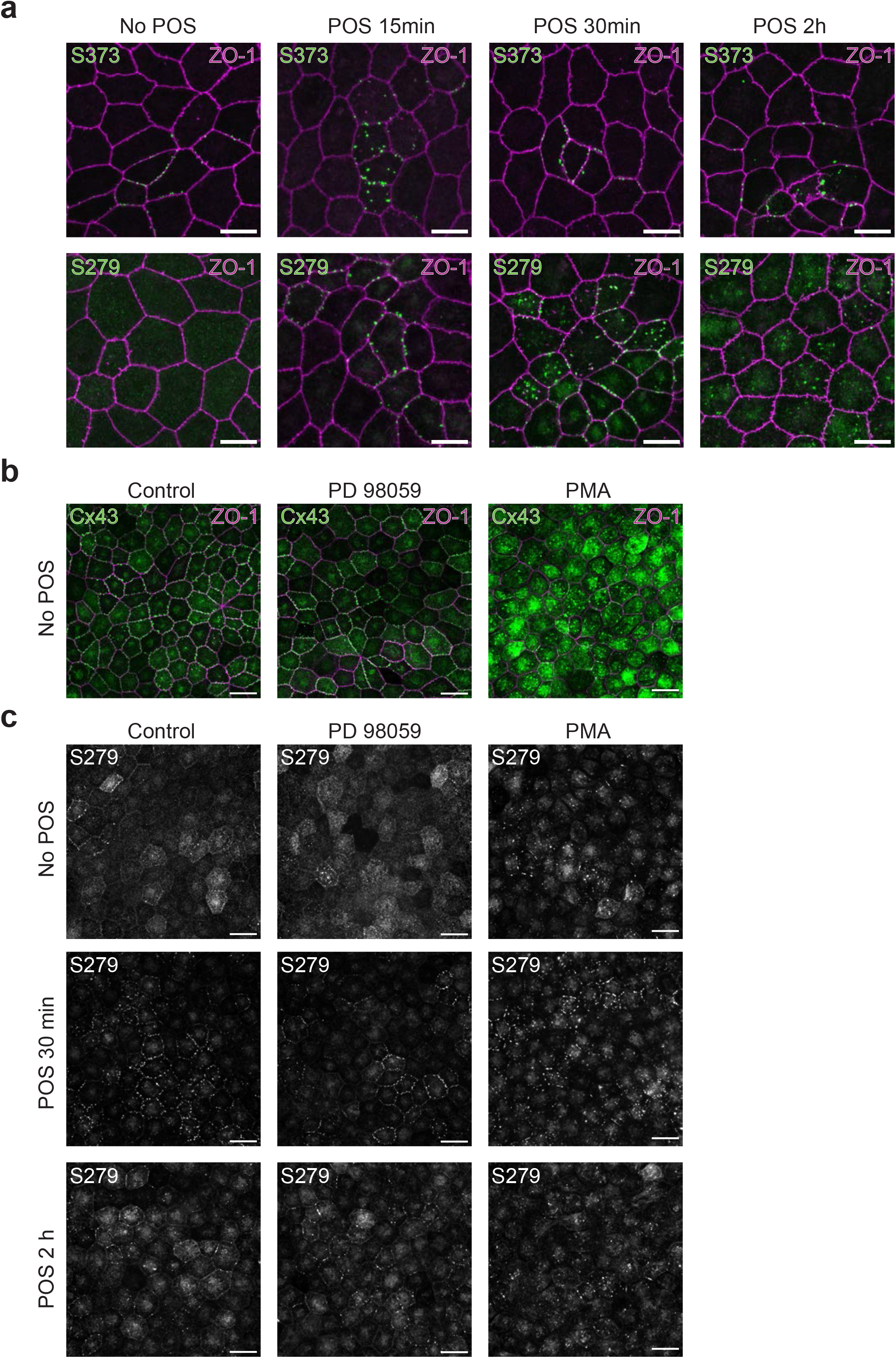
Gap junctions are phosphorylated at S279 during phagocytosis and PKC activation induces gap junction dissociation **a**. Serine 373 (S373, top) and serine 279 (S279, bottom) phosphorylated forms of Cx43 (*green*) and ZO-1 (*magenta*) immunostaining with no POS as well as 15 min, 30 min, and 2 hr post POS application in hESC-RPE. Scalebars 10 µm. **b**. Effect of vehicle (left), MEK inhibition (PD 98059, middle), and PKC activation (PMA, right) on Cx43 (*green*) and ZO-1 (*magenta*) staining. Scalebars 15 µm. **c**. Same as **b** except staining for the Cx43 phosphorylated at S279 during the phagocytosis challenge. Scalebars 15 µm.

**Figure 4:**
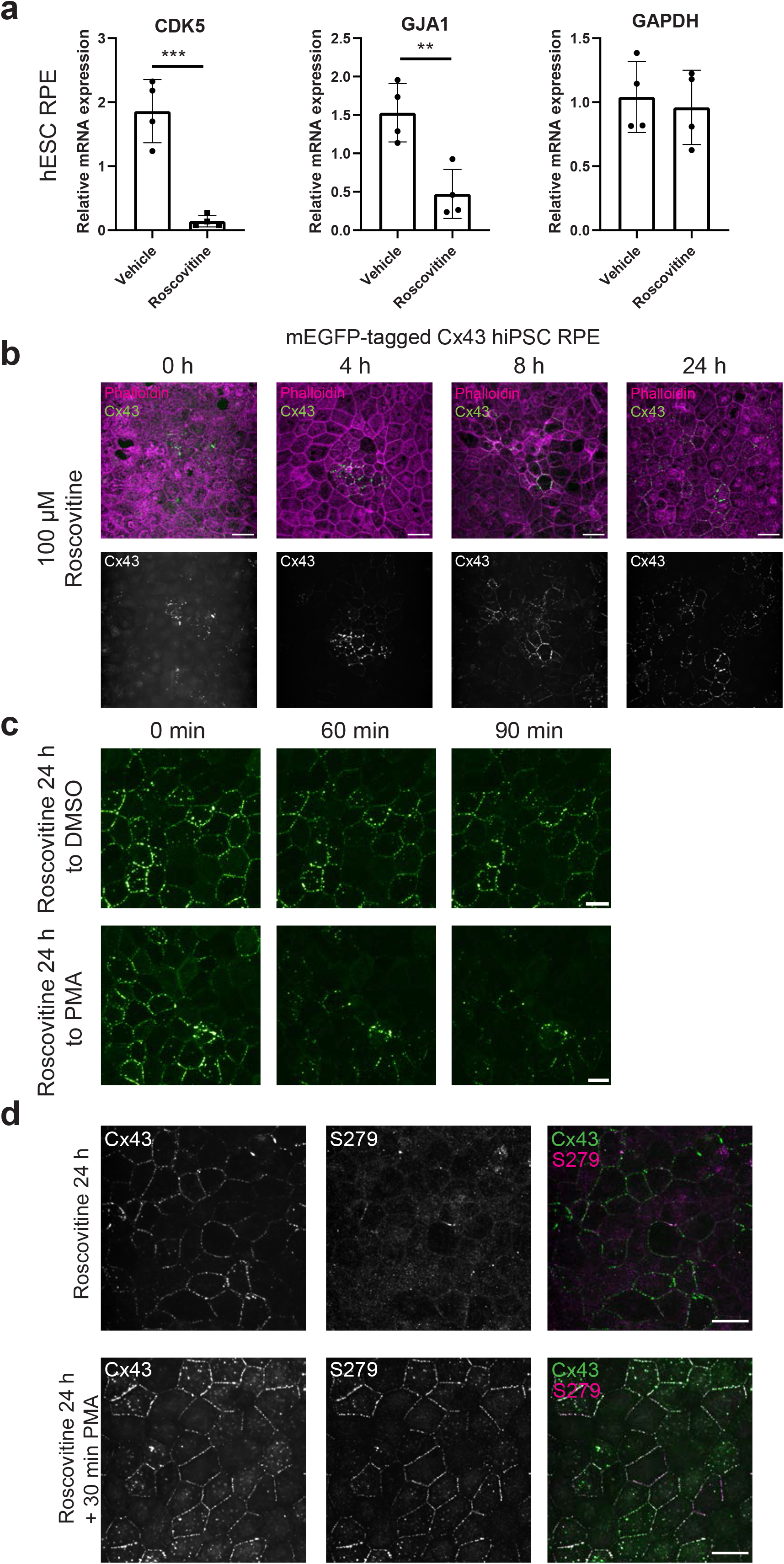
Effects of a CDK5 inhibitor on Cx43 localization and regulation **a**. mRNA expression levels for CDK5, GJA1 (Cx43), and GAPDH in vehicle and roscovitine treated hESC-RPE (n = 4 experiments). **b**. Cx43 (*green*, bottom) and phalloidin (*magenta*) labeling in the mEGFP-tagged Cx43 hiPSC-RPE at different time points after treatment with roscovitine, a Cdk5 inhibitor. Scalebars 15 µm. **c**. Maximum intensity z-projections from Z-stacks captured during live imaging of mEGFP-tagged Cx43 hiPSC-RPE showing the effect of reversal of a 24 hr roscovitine treatment on Cx43 (*green*) localization switching to either a DMSO (top) or PMA (bottom) solution over time. Scalebars 10 µm. **d**. Maximum intensity z-projections from mEGFP-tagged Cx43 hiPSC-RPE (*green*) after 24 h roscovitine treatment with or without subsequent PMA incubation for 30 min labeled for S279 phosphorylated form (*magenta*). Scalebars 15 µm.

We focused our further investigation on MAPK and PKC signaling cascades, as they both have previously been shown to phosphorylate Cx43 and to be necessary for the disruption of Cx43 activity (Hossain et al., 1998; Lampe and Lau, 2004). Compared to vehicle-treated cells (Figure 3b), inhibiting MEK (PD 98059), an upstream kinase in the MAPK cascade (Nimlamool et al., 2015), did not result in any significant difference in general Cx43 labeling. However, treating the cells with a PKC activator (PMA) caused a dramatic translocation of Cx43. This effect was more potent than was observed to occur during phagocytosis (Figure 4b, Figure 2a). In both experiments, the cells were incubated for 2.5 h to reflect the timescale of phagocytosis experiments. Since the prominence of S279 phosphorylation was affected by phagocytosis, we wanted to investigate the effects of blocking MAPK or activating PKC during phagocytosis (Figure 3c). Here, the samples were preincubated with the kinase modulators or DMSO for 30 min before the addition of POS particles. We found no change in the level of S279 labeling between DMSO control and PD 98059 in the no POS samples. Furthermore, inhibiting MAPK kinase did not prevent the emergence of S279 labeling that we had observed during POS phagocytosis. As expected, activating PKC did not prevent the phagocytosis evoked phosphorylation either, but it accelerated the translocation as the labeling pattern was found much more diffuse already at 30 min when compared to the vehicle-treated phagocytosis samples (Figure 3c). By activating PKC, labeling for S279 was already detected without the addition of POS particles. Therefore, we concluded that phosphorylation of S279 regulated gap junction internalization, and that a potential effector was PKC.

### The level of junctional Cx43 is regulated by cyclin dependent kinase 5

In addition to interacting with MAPK and PKC pathways, Cdk5 kinase has been shown to directly phosphorylate Cx43 at S279 during embryonic development (Qi et al., 2016). We wanted to investigate whether this kinase was present in mature RPE, and how inhibiting its activity with roscovitine modulated the levels of Cx43. In order to investigate whether Cdk5 could be present in mature RPE, we decided to investigate mRNA levels first (Supplemental Figure 1). The levels were quantified from hESC-RPE that had either been treated with roscovitine or equivalent amount of DMSO (vehicle) for 24 h (Figure 4a, Supplementary Figure 1). Our results showed that Cdk5 was indeed found in RPE, and that its expression significantly decreased after the inhibitor treatment (p-value = 0.0005). Furthermore, the mRNA level for Cx43 (GJA1) was significantly reduced in the roscovitine treated cells (p-value = 0.0053). Levels of GAPDH were not changed between control and roscovitine samples (p-value = 0.7022) (n = 4 for all groups).

We investigated the effect of roscovitine on Cx43 at the protein level by imaging the EGFP-Cx43 expressing hiPSC-RPE cells at 0 h, 4 h, 8 h and 24 h time points post roscovitine application (Figure 4b). Because we were imaging a monoallelic cell line, at 0 h we often found areas with limited junctional Cx43 foci. However, with the roscovitine treatment, the junctional foci increased with each timepoint and by 24 h, most of the junctions were positive for Cx43. Next, we wanted to analyze whether roscovitine treatment could be reversed by replacing the roscovitine supplemented medium after 24 h either with control medium (DMSO) or PMA, and imaging the cells live for 90 min. The obtained Z-stacks showed that removing roscovitine alone did not cause the Cx43 to translocate, but with the addition of PMA, a substantial portion of the foci had disappeared after 90 min (Figure 4c). These results suggest that PMA-induced activation of PKC causes a similar translocation that we observed in phagocytosis.

After demonstrating that roscovitine increased junctional Cx43, we wanted to investigate whether inhibiting Cdk5 kinase also had an effect on S279 phosphorylation (Figure 4d). Our immunolabeling experiments indicated that incubating the EGFP-Cx43 expressing hiPSC-RPE with roscovitine alone for 24 h did not induce the S279 phosphorylation. Yet, when the 24 h incubation was followed with 30 min incubation with PMA, most of the foci were also positive for S279. This indicates that the S279 phosphorylation occurs before the gap junctions are dissociated.

### Inhibiting Cdk5 increases gap junctional conductance and decreases phagocytosis efficiency

As Cdk5 inhibition increases junctional labeling of Cx43, the natural question is whether it has functional consequences on the resulting gap junctional conductances (Figure 5a). Simultaneous paired patch clamp recordings were performed from hESC-RPE monolayers that were either treated with DMSO (Control) or roscovitine for 24 h. The cells were kept in the recording chamber for a maximal time of 90 min as we had seen that the roscovitine effect would not be significantly reversed in this timespan (Figure 4c). Our resulting current clamp recordings showed that roscovitine treated hESC-RPE cells had significantly higher levels of gap junctional coupling when compared to control (p-value = 0.0020, n = 5 pairs for each condition).

**Figure 5:**
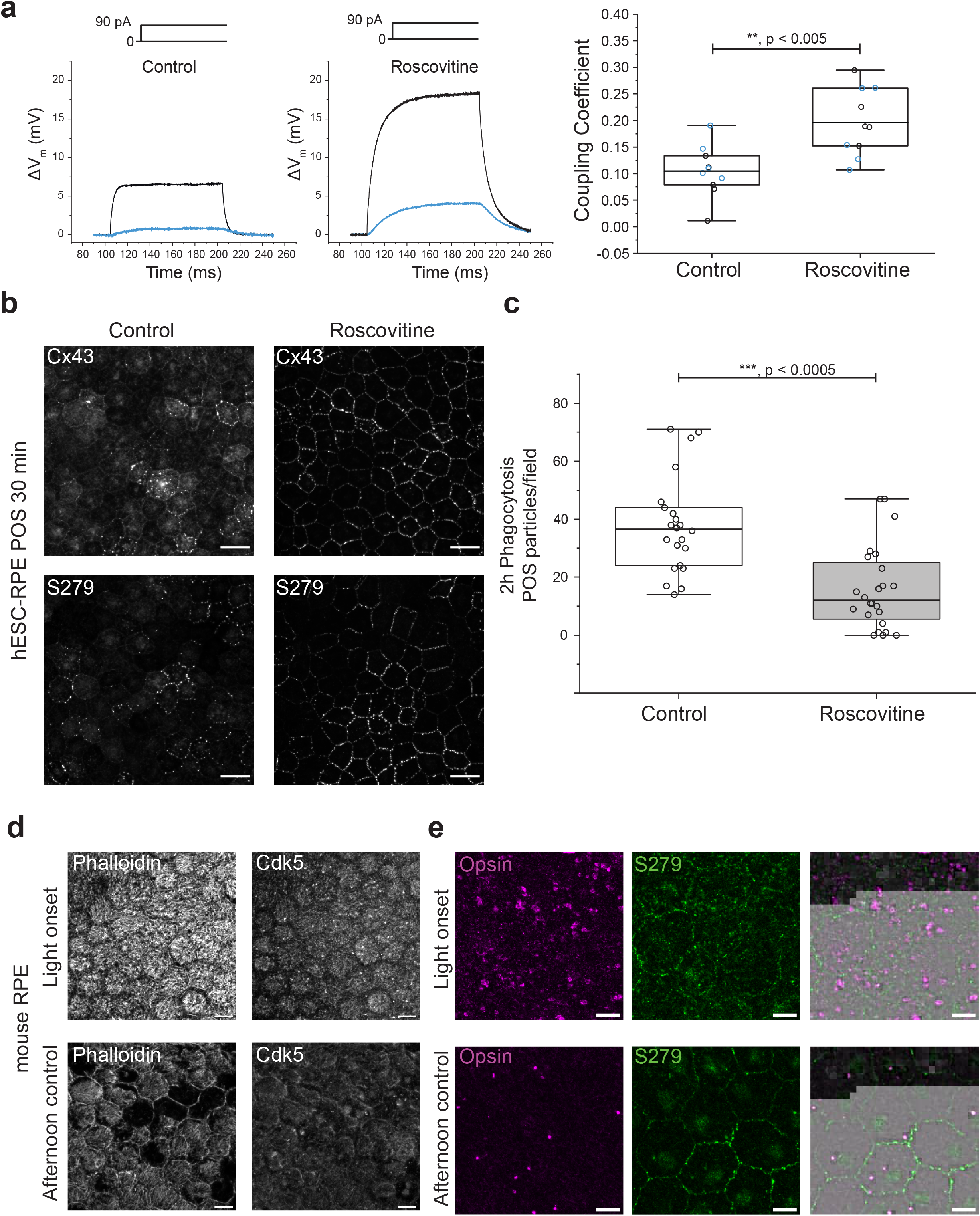
Circadian regulated Cdk5 impacts gap junctional communication and has an effect on phagocytosis efficiency **a**. Coupling coefficient analysis from paired patch clamp recordings of hESC-RPE with and without 24h roscovitine treatment. Example traces (left) show current injected into cell 1 (*black*) and the response in cell 2 (*blue*), and the population data is shown on the right (n = 5 pairs). Center lines show the means (Statistical analysis with Student’s t test); box limits indicate the 25th and 75th percentiles; data points are plotted as open circles where *black* indicates coupling from cell 1 to cell 2 and *blue* is coupling from cell 2 to cell 1. **b**. Cx43 (top) and S279 (bottom) localization in hESC-RPE 30 min after POS treatment in control conditions and after 24h roscovitine treatment (right). Scalebar 15 µm. **c**. Quantification of POS particles after 2 h phagocytosis assay in control and roscovitine samples (**b**) (n = 3 experiments). Center lines show the medians (Statistical analysis with Mann-Whitney U Test); box limits indicate the 25th and 75th percentiles; data points are plotted as open circles. **d**. Phalloidin (left) and Cdk5 (right) labeling in mouse RPE monolayers at light onset (top) and afternoon (bottom). Scalebar 15 µm. **e**. Opsin (*magenta*) and S279 (*green*) labeling in mouse RPE monolayers at light onset (top) and afternoon (bottom). Scalebar 10 µm.

Next, we wanted to analyze whether treating hESC-RPE with roscovitine for 24 h affected the translocation that we had observed during phagocytosis (Figure 5b). We focused this analysis on the 30 min time point as that had shown to be most significant for the S279 phosphorylation (Figure 4). In vehicle-treated hESC-RPE, S279 signal was evident, and the sample showed efficient translocation of Cx43. Interestingly, this translocation was significantly inhibited in the roscovitine treated cells, and much of the Cx43 and S279 labeling was still present in the junctional area in the hESC-monolayers. To investigate whether this diminished translocation had also an effect on the phagocytosis efficiency, we quantified the total number of POS particles after 2 h incubation either in DMSO (control) or roscovitine treated hESC-RPE samples (Figure 5c). Both groups received 24 h preincubation prior to the onset of phagocytosis. By quantifying the number of POS particles, we found the phagocytosis rate significantly decreased in the roscovitine treated group (p-value = 0.000054, n = 3 inserts for each group). Taken together, this data indicated that CDK5 activation is necessary for relocalization of Cx43 away from junctions which in turn is essential for phagocytosis.

We investigated whether Cdk5 was also present in mouse RPE *in vivo* by immunolabeling at different points of the diurnal cycle (Figure 5d,e). The results showed that at light onset when phagocytosis occurred in bulk, the kinase localized into specific clusters at the plasma membrane. Instead, in the samples that had been fixed in the afternoon outside phagocytosis peak, the labeling of Cdk5 was found more diffuse. Different from hESC-RPE, in mouse RPE, S279 was identified at both timepoints (Figure 6d), but the apical membrane labeling was much more prominent at light onset whereas in the afternoon the localization was more concentrated into specific foci. Moreover, S279 was found to localize adjacent to opsin. Opsin labeling was dramatically lower in the afternoon when compared to light onset, which is expected considering that POS phagocytosis peaks in the morning (LaVail, 1976).

**Figure 6:**
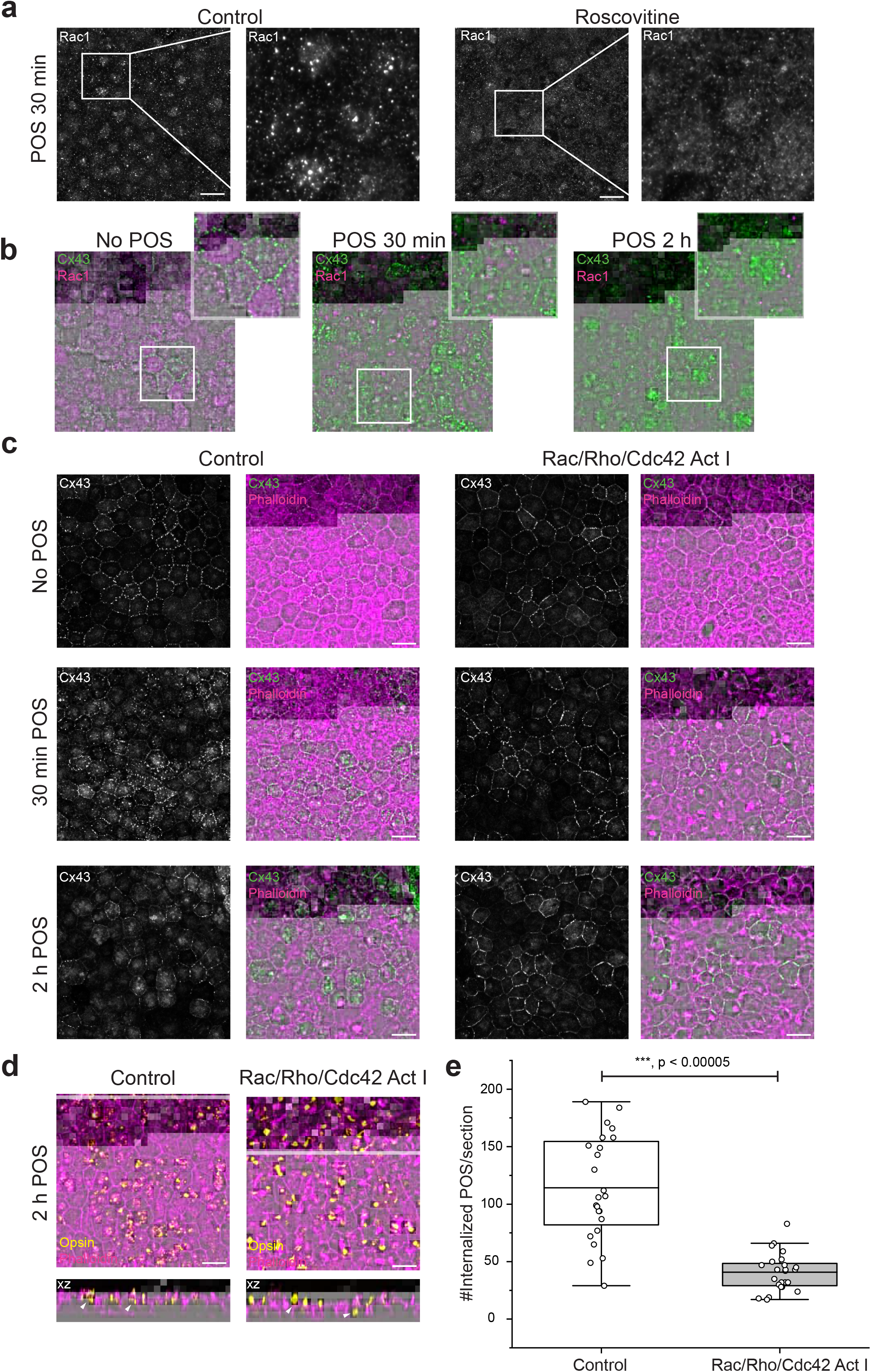
Rac1 pathway activation regulates maintenance of Cx43 at the junction **a**. Rac1 labeling 30 min after POS particle application in control conditions or after 24 hr incubation with roscovitine. **b**. Cx43 (*green*) and Rac1 (*magenta*) labeling with no POS, 30 min post POS application, and 2 hr post POS application. Scalebars 15 µm. **c**. Cx43 (*green*) and F-actin (phalloidin, *magenta*) labeling at the three time points used above. Both in control conditions (left) and in the presence of Rac/Rho/Cdc42 Activator I (right). Scalebars 15 µm. **d**. Opsin (I) and F-actin (phalloidin, *magenta*) labeling at phagocytosis timepoints as above in control conditions (left) and in the presence of Rac/Rho/Cdc42 Activator I (right). Scalebars 15 µm. **e**. Quantification of POS particles (see (Viheriälä et al., 2021) for analysis details) that have been internalized by hESC-RPE after 2 h phagocytosis assay in control and Rac/Rho/Cdc42 Activator I treated conditions (n = 4 experiments). Center lines show the means (difference analyzed with Student’s t test); box limits indicate the 25th and 75th percentiles; data points are plotted as open circles.

### Rac1 activation reduces Cx43 internalization

To better understand the regulatory role of Cdk5 in gap junction dynamics during POS phagocytosis, we wanted to investigate possible downstream mechanisms of this pathway. Interestingly, Cdk5 has previously been shown to regulate actin dynamics during synaptic plasticity (Fu et al., 2007) and neuronal migration, and most of these regulator roles have been linked to the family of Rho GTPases (Shah and Rossie, 2018). The formation of phagocytic cups requires dramatic reorganization of the actin cytoskeleton, and the recruitment of F-actin to the forming cups has been shown to involve the activation and relocalization of a specific Rho GTPase, Rac1 (Mao and Finnemann, 2012). Moreover, Cdk5 has been shown to suppress Rac1 activity during cellular senescence (Alexander et al., 2004). Considering all this, we studied the localization of Rac1 during the POS challenge with and without roscovitine in hESC RPE (Figure 6a). Our results showed that in control conditions, Rac1 aggregated into specific clusters on the apical membrane as Mao and Finnemann (2012) had shown previously, but this labeling pattern was not as evident in roscovitine treated samples implicating the activation of Rac1 could be affected.

As the apical clusters of Rac1 resembled the pattern we had observed for Cx43 during phagocytosis, we wanted to co-label Rac1 and Cx43 during different stages of phagocytosis. Our immunolabeling showed that in control conditions (no POS), Rac1 labeling was very diffuse, and no co-localization was found with the junctional Cx43. In contrast, after 30 min incubation some, but not all, of the apical Rac1 clusters were also positive for Cx43. After 2 hours, the aggregation of Rac1 was no longer detected. Therefore, similar to how Cx43 shows temporal dynamics in response to phagocytosis, Rac1 also changes its cellular distribution.

Given that Rac1 demonstrated temporal dynamics in response to phagocytosis, our next step was to investigate how direct Rac1 activation would affect the localization and dynamics of Cx43 during phagocytosis. To test this, we used Rac/Rho/Cdc42 Activator I (Act I; Figure 6c) and looked at the same phagocytosis time points as before. Since Rac1 and its downstream partners are known to affect the actin cytoskeleton, we performed phalloidin staining in addition to staining for Cx43 to examine its effects. While the translocation of Cx43 was detected in the vehicle-treated control samples, it was found to remain in the junctions in the Activator I-treated samples (Figure 6c). In fact, after 2 h POS incubation, the Activator I samples had stronger junctional Cx43 labeling than the no phagocytosis group. This stronger junctional labeling also resulted in a decrease in Cx43 at the mRNA level (p-value = 0.0175) while the levels of GAPDH were not changed between Activator I and control samples (p-value = 0.9914) (n = 3 for all groups), similarly to what we had observed in roscovitine treated samples (Supplemental Figure 1 c,d). Furthermore, we observed higher recruitment of F-actin to nascent phagosomes in the Activator I samples when compared to the control phagocytosis samples. The number of internalized POS was also found to be significantly reduced in the presence of Activator I (Figure 6d,e; p = 3 × 10^−8^). Therefore, as continuous activation of Rac1 decreased the translocation of Cx43 connexons during phagocytosis, it appeared to inhibit the internalization of POS particles, similarly to Cdk5 inhibition.

## Discussion

Dynamic changes in gap junctional coupling and Cx hemichannel activity have long been recognized as important regulators of embryonic brain development and cell division (Levin, 2007; Sutor and Hagerty, 2005). Synchronizing information flow within tissue and having electrically isolated embryonic cells are both known to be critical to ensure properly regulated morphogenesis (Aslanidi et al., 1991; Rela and Szczupak, 2004; Sutor and Hagerty, 2005). Gap junctions have been shown to reconfigure microcircuits of the neural retina on both a millisecond timescale as well as over the course of the circadian cycle (Krizaj et al., 1998; Li et al., 2013; Ribelayga et al., 2008; Trenholm and Awatramani, 2019; Völgyi et al., 2013). Since the retina and RPE are intimately interlinked in the eye, it is likely that similar dynamics could occur in RPE.

RPE exhibits an extensive network of Cx43 proteins. While the electrical coupling had turned out to be surprisingly low, it was readily modifiable (Fadjukov et al., 2022). Therefore, we asked whether gap junctions could serve a role in one of the most time-restricted tasks of the mature RPE: phagocytosis of photoreceptor outer segments. The reported timescale of the phagocytosis in vivo (Bosch et al., 1993; LaVail, 1976) coincides with the half-live for gap junctions (Laird et al., 1991; Rhett et al., 2011). In addition, various voltage-gated ion channels have been previously reported to contribute to the pathway (Johansson et al., 2019; Korkka et al., 2019; Mamaeva et al., 2021; Müller et al., 2014), and thus gap junctions could play an important role in regulating electrical properties of RPE.

In this study, we discovered that Cx43 undergoes a dramatic translocation during the phagocytosis process, and the timescale of their turnover follows that reported for gap junction disassembly that occurs in response to injury or growth factors (Lastwika et al., 2019; Nimlamool et al., 2015; Solan and Lampe, 2014). After 30 min of POS challenge, the number of gap junctions appeared to decrease being in various stages of internalization, and most of the junctional labeling had disappeared after 2 h. Interestingly, it has been shown that ZO-1 could regulate the transition of undocked connexons into gap junctions, while disruption of Cx43/ZO-1 interaction could enlarge the size of gap junctions (Hunter et al., 2005; Rhett et al., 2011). It is worth noting that our electrophysiological recordings did not on average show significant changes in input resistance in hESC-RPE that had been treated with POS (Supplemental Figure 2a). We also recorded from mouse RPE dissected shortly before to light onset (Supplemental Figure 2b) and occasionally observed a cell with especially high input resistance. However, the formation of phagocytic cups can change the membrane properties and as a result could mask the possible effects in input resistance. Moreover, any potential changes in gap junctional conductance are probably fast and might not occur in every cell, thus these changes would be difficult to detect in experimental conditions. In addition, Cx43 hemichannels are known to open in response to integrin α5 and shear stress (Leithe et al., 2018) which could counteract the closure of Cx43 gap junctions in the cell-cell junctions. In one example cell (Supplemental Figure 2c) we observed that the input resistance was drastically reduced from high to control levels during the time course of the patch. Thus, while gap junctions are temporally and spatially regulated during POS phagocytosis, it may be difficult to directly measure the accompanying changes in membrane properties.

At this stage it is not certain whether it is only the Cx C-terminus or the entire gap junction that undergoes disassembly during the phagocytosis process as all our means to visualize gap junctions target that part of the protein structure. It is known that the C-terminal end can serve channel-independent roles and act as a signaling scaffold for various kinases to interact with the cytoskeleton and play a role e.g. in cell migration (Kameritsch et al., 2012; Leithe et al., 2018; Solan and Lampe, 2018). Interestingly, the membrane translocation of Rac1 has previously been shown to be controlled by integrins and it is believed to be critical for the activation of Rac effector Pak (Del Pozo et al., 2002). Studies on filopodia formation have demonstrated that interaction of Pak and Cx43 enables the effector to be more activated and allows a stronger phosphorylation of its downstream targets p38 and Hsp27 to promote migration (Kameritsch et al., 2015). It is possible that phagocytic cup formation requires similar interactions in RPE as it also involves a reorganization of the actin cytoskeleton. Interestingly, the activation of Rac1 requires spatiotemporal regulation and closure of the phagocytic cup is delayed in cells that express constitutively active Rac1 (Nakaya et al., 2008). Moreover, in RPE cells inhibiting the activity of Rac1 results in the lack of F-actin assembly and reduction in the number of engulfed POS particles (Mao and Finnemann, 2012). This is in line with our results as we detected Cx43 and Rac1 in the same clusters only periodically and observed an increase in F-actin recruitment. Constitutively active Rac1 also seemed to delay the translocation of Cx43. The quantification of internalized POS particles showed that constitutively activating Rac1 and delaying Cx43 translocation decreased the efficiency of phagocytosis but did not fully prevent POS internalization. Moreover, the images indicate that POS are more readily fragmented in control conditions, while in Rac1 treated samples large particles are visible inside the cells. The implication of Cx43 in the Rac1 pathway is also interesting as the activation of Rac1 is thought to be voltage-dependent (Yang et al., 2020), and Cx43 strongly colocalized with Na_V_1.4 that also changed its localization during the phagocytosis pathway (Johansson et al., 2019).

The regulation of Cx43 by phosphorylation has been extensively studied (Hossain et al., 1998; Lampe and Lau, 2004; Solan and Lampe, 2014, 2018), and we focused our investigation on the residues and kinases that have previously been highlighted in gap junction turnover. Of the phosphorylated residues, S373, while not dynamic throughout the course of phagocytosis in our study, could explain why the connectivity has been shown to vary between the different cells in the RPE monolayer (Fadjukov et al., 2022). S373 phosphorylation has been shown to serve as a molecular switch to rapidly increase gap junctional communication (Dunn and Lampe, 2014). Thus, the paucity lack of this residue could also explain why input resistance levels were not consistently significantly different in RPE between night and day (Supplemental Figure 2b). It could also highlight the difference between the constant requirement of phagocytosis of RPE to acute injury that requires more immediate changes in cellular connectivity. In the cases with chronic disassembly signal, including when v-src is chronically coexpressed with Cx43, this kinase can bypass the need for S373 modulation by Akt kinase by directly displacing ZO-1 from binding to Cx43 (Solan and Lampe, 2018; Sorgen et al., 2004; Toyofuku et al., 2001).

Our study showed that S279 was the most significant modulation during POS phagocytosis. The difference between control and phagocytosis conditions were more drastic in hESC-RPE than mouse RPE, but it is important to note that while the burst of phagocytosis is specific in vivo, there remains a baseline level of POS degradation throughout the day (Ko et al., 2009; Nandrot et al., 2004; Strauss, 2005). It is also possible that S279 could serve as a mechanism to prime the epithelium for the burst. In addition, it is likely that gap junctions are even more dynamic in tissue, with new hemichannels continuously added to replace the internalized gap junctions. Cx43 phosphorylation events can occur within 15 min of gap junction synthesis (Solan and Lampe, 2018). The lack of this phosphorylation in non-phagocytosis conditions in cultured RPE, and thus the lack of proposed priming, could partially explain why cultured cells tend to have a slower time course of phagocytosis (Mazzoni et al., 2014). On the other hand, the constant presence of this phosphorylation in mouse RPE cells could explain why these cells have low levels of coupling as S279 has been shown to induce reduction of channel open probability and accordingly decreased gap junction communication (Cottrell et al., 2003).

Our data indicates that Cx43 translocation requires sequential activity of multiple kinases, and that inhibiting the MAPK pathway alone was not sufficient to prevent the emergence of S279. It is possible that activity of v-Src promotes S279 phosphorylation as has been shown by others (Solan and Lampe, 2008, 2014). As this kinase was not inhibited in this study, and we did not investigate its target tyrosine phosphorylation sites Y247 and Y265 (Solan and Lampe, 2008), the effect of this kinase cannot be ruled out. Studies in other phagocytes such as macrophages have shown that phagosome formation requires changes in the membrane lipid composition mediated by phospholipase Cγ that in turn would lead to PKC activation via diacylglycerol production (Uribe-Querol and Rosales, 2020). In this study, we did not investigate the effect of PKC modulation on POS phagocytosis efficiency, but earlier studies have found that while PMA did not alter particle binding, it reduced internalization as did inhibition of PKC (Finnemann and Rodriguez-Boulan, 1999; Hall et al., 1993). PKC activation and synthesis in the retina are known to be regulated by diurnal rhythm (Gabriel et al., 2001). PKC phosphorylation has also been shown to proceed MAPK phosphorylation in response to vascular vascular endothelial growth factor and other stimuli (Nimlamool et al., 2015; Sorgen et al., 2018). This is in line with our experimental results and could explain why S279 levels were higher after PMA treatment. Moreover, 12-O-tetradeconoylphorbol-13-acetate, another pharmacological inducer for PKC, has been shown to promote gap junction disassembly, similarly that we observed with PMA (Lampe, 1994). At the time of this study, we did not have an antibody for S368, the most recognized target for PKC (Lampe et al., 2000), that would be suitable for immunocytochemistry.

In addition to MAPK and PKC, Cx43 has been shown to be modulated by Cdk5 that is primarily active in postmitotic neurons (Contreras-Vallejos et al., 2012; Liu et al., 2008; Qi et al., 2016). Roscovitine treatment in RPE dramatically increased junctional Cx43, while its mRNA levels were found to decrease. Inhibiting Cdk5 could deplete the cytoplasmic pool of connexons while the formed junctions are targeted towards the plasma membrane. Previous work has shown that Cdk5 can inhibit the membrane targeting of Cx43 (Qi et al., 2016). In addition, Cdk5 has been shown to directly induce the S279 phosphorylation (Qi et al., 2016), but we found that it can occur even after roscovitine treatment. It is possible that residual levels of the activity are sufficient to induce the phosphorylation, or it could be due to the activity of other kinases activated by PKC. The dramatic effect of PMA could be explained by the fact that PKC has been suggested to activate the Cdk5/p25 complex (de Thonel et al., 2010). While PMA was found to induce Cx43 internalization even without POS exposure, physiological levels of PKC activation during phagocytosis were not sufficient to induce translocation in roscovitine treated samples. These results suggest that PKC and Cdk5 are both required to regulate Cx43 translocation during POS phagocytosis.

Because the Cdk5 inhibition decreased POS phagocytosis efficiency, the translocation of Cx43 is most likely crucial for the phagocytosis process. Interestingly, Cdk5 has also been recently shown to regulate the circadian clock, and its silencing in the suprachiasmatic nucleus shortens the clock period (Brenna et al., 2019; Kwak et al., 2013). While Cdk5 protein levels were not found to change, its activity was highest during the night and lowest during the day (Brenna et al., 2019). Moreover, roscovitine treatment in striatum neurons has been shown to evoke dopamine release (Chergui et al., 2004). Dopamine levels in the eye vary according to circadian rhythm and have been shown to affect the coupling of rods and cones (Goel and Mangel, 2021; Ribelayga et al., 2008). RPE has also been shown to have a circadian clock that is entrained by dopaminergic signaling, while removal of its dopamine receptors can prevent the phagocytosis burst (Baba et al., 2017; Goyal et al., 2020). The C57BL/6 mice used in this study lack a circadian rhythm in their dopamine levels if kept in constant darkness without cyclic light (Dinet et al., 2007; Korshunov et al., 2017). The burst of phagocytosis still occurs at light onset, thus the phagocytic process is still maintained (Vargas and Finnemann, 2022). We propose that Cdk5 could play a role in maintaining the circadian timing of phagocytosis in RPE. The diurnal nature of Cx43 translocation could explain why its role in phagocytosis is more definitive in RPE than in macrophages (Anand et al., 2008; Dosch et al., 2019; Glass et al., 2013). Our immunostainings also suggest differences in Cdk5 activity between night and day, and this spatiotemporal change, with the cooperation from other kinases, could help to initiate the translocation of Cx43 in RPE.

The role of Cdk5 in POS phagocytosis is strengthened by the fact that this kinase has previously been shown to regulate endocytosis of synaptic vesicles (Tan et al., 2003; Tomizawa et al., 2003). Lastly, Cdk5 has been implicated in Rac1 regulation in cellular senescence that could explain why we observed a similar Cx43 translocation deficiency with both roscovitine and Rac1 activator. Further studies are needed to fully elucidate the roles that Cdk5 and Cx43 play in phagocytosis in the mature RPE, but our data demonstrate that both are important for regulating this important task (a proposed pathway schematic in Figure 7). Translocation of gap junctions during phagocytosis could enable RPE cells to electrically decouple, facilitating the activation of voltage-gated channels that have been previously shown to play a role in phagocytosis (Fadjukov et al., 2022). Ultimately, the critical role of gap junctions in phagocytosis could help to better understand the pathological defects of this pathway.

**Figure 7:**
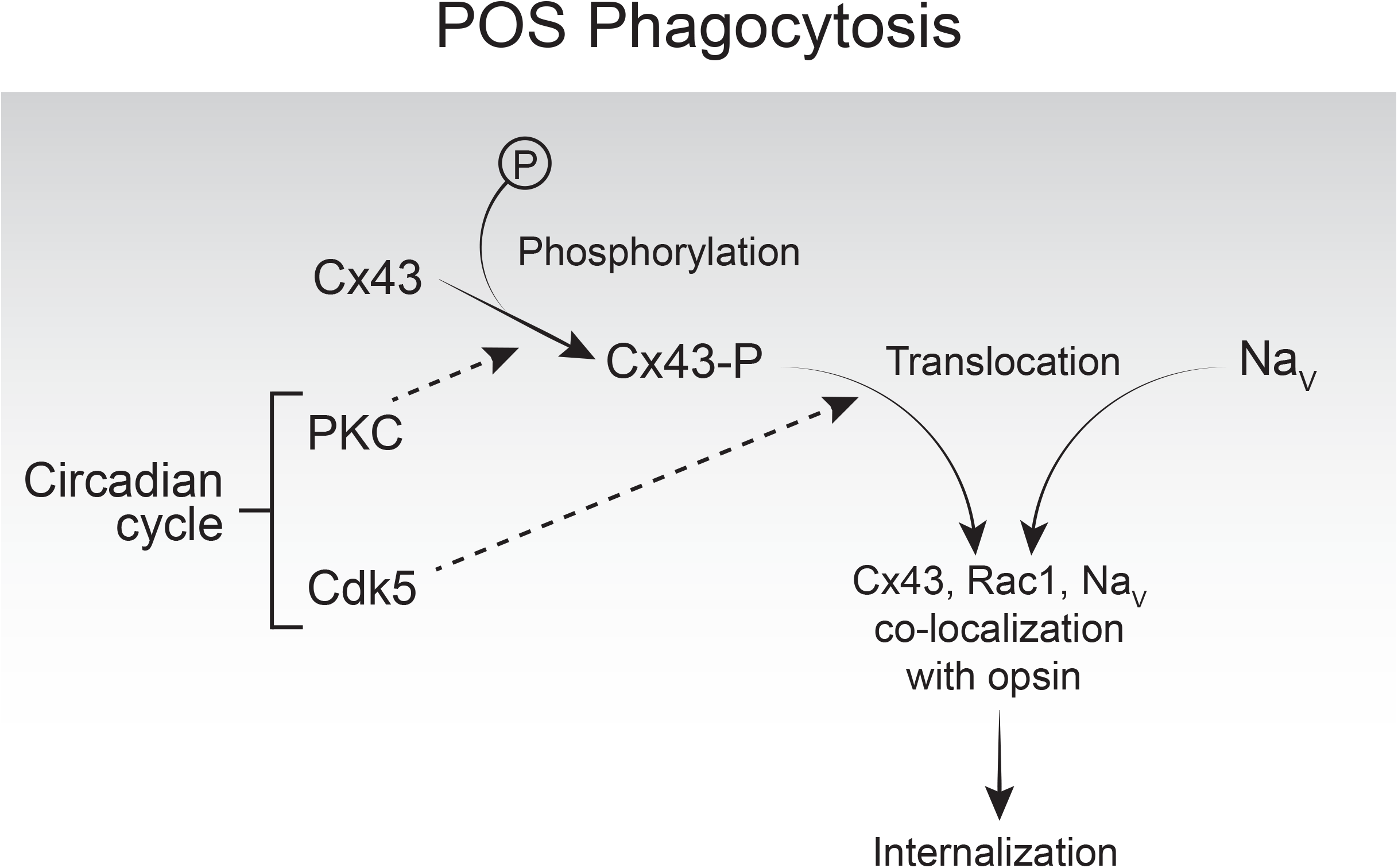
A proposed pathway for the mechanism for the role of Cx43 in POS phagocytosis. In control conditions, Cx43 strongly co-colocalizes with NaV1.4. Junctional Cx43 are phosphorylated by kinases such as PKC whose activity is regulated by the circadian rhythm. During phagocytosis, the translocation of Cx43 is regulated by Cdk5 that can also regulate the circadian clock and suppress the activity of Rac1. Once Cx43 are moved to the apical membrane, they co-localize with Rac1, opsin and NaV. As the phagocytosis pathway progresses, Cx43 are internalized with the phagosomes.

## Acknowledgements

We would like to acknowledge Outi Heikkilä (Tampere University) and Susan Lynn Wohlgenant (Northwestern University) for their expertise and assistance, and Viivi Karema-Jokinen (Tampere University) for her technical assistance with the electron microscopy experiments. We are grateful to Dr. Heli Skottmann (Tampere University) for kindly providing the hESCs for RPE differentiation and for Dr. Keijo Viiri (Tampere University) for the availability of mouse tissue. The authors acknowledge the following cores: Biocenter Finland (BF), Tampere Imaging Facility (TIF), and Northwestern Center for Advanced Microscopy (CAM). In addition, Tampere Facility of Electrophysiological Measurements, and Electron Microscopy Unit (University of Helsinki, Institute of Biotechnology) are gratefully acknowledged for their services.

This study was funded by the Academy of Finland (grant numbers 319257, 287287, 323507, 308315, 340127 and 330896), the Emil Aaltonen Foundation, the Jane and Aatos Erkko Foundation, TUT on world tour mobility grant (Tampere University), the Finnish cultural Foundation, Oskar Öflunds Foundation, Ella and Georg Ehrnrooth Foundation, Federation of European Biochemical Societies (FEBS) Short-Term Fellowship.

## Author Contributions

J.F., S.W., G.W.S., and S. N. contributed to experimental design as well as data acquisition, analysis, and interpretation. N.M. performed the mRNA quantification and analysis. S.H. and M.V.R. planned and conducted the electron microscopy experiments including sample preparation (with J.F.) and imaging. T.I. contributed expertise on confocal microscopy and image analysis. All authors contributed to the writing of the manuscript and approved the final manuscript.

## Supplemental Materials

**Supplemental Figure 1:** mRNA quantification for CDK5, GAPDH and GJA1.

**a**. Table showing details of the primers and PCR conditions that were used to quantify mRNA levels for CDK5 and GJA1, GAPDH was used as the reference gene.

**b**. Example of the PCR products electrophorized on gel that were used for band quantification. First four lanes correspond to roscovitine treated samples, and the last four lanes are vehicle (DMSO) treated samples.

**c**. mRNA expression levels for GJA1 (Cx43), and GAPDH in vehicle and Rac/Rho/Cdc42 Activator I treated hESC-RPE (n = 3 experiments).

**d**. Example of the PCR products that were used for band quantification. First three lanes correspond to vehicle (cell culture medium), and the last four lanes are Rac/Rho/Cdc42 Activator I treated samples.

**Supplemental Figure 2:** Changes in RPE input resistance with phagocytosis

**a**. Input resistance analysis of hESC-RPE cells current clamp recordings in control conditions and after 30 min or 2 h POS challenge. Lines correspond to mean values.

**b**. Input resistance analysis of mouse RPE cell current clamp recordings that were performed at various points of the circadian cycle (n = 3 animals, 146 cells).

**c**. One example mouse cell from (**b**) that had a noticeable change in input resistance throughout the duration of the recording. Time 0 was the start of the recording; the input resistance is noted above each trial.

## References

Alexander, K., Yang, H.-S., and Hinds, P.W. (2004). Cellular senescence requires CDK5 repression of Rac1 activity. Mol. Cell. Biol. 24, 2808–2819..

Anand, R.J., Dai, S., Gribar, S.C., Richardson, W., Kohler, J.W., Hoffman, R.A., Branca, M.F., Li, J., Shi, X.-H., Sodhi, C.P., et al. (2008). A role for connexin43 in macrophage phagocytosis and host survival after bacterial peritoneal infection. J. Immunol. 181, 8534–8543..

Aslanidi, K.B., Boitsova LJu, Chailakhyan, L.M., Kublik, L.N., Marachova, I.I., Potapova, T.V., and Vinogradova, T.A. (1991). Energetic cooperation via ion-permeable junctions in mixed animal cell cultures. FEBS Lett. 283, 295–297..

Baba, K., DeBruyne, J.P., and Tosini, G. (2017). Dopamine 2 Receptor Activation Entrains Circadian Clocks in Mouse Retinal Pigment Epithelium. Sci. Rep. 7, 5103..

Bobu, C., Craft, C.M., Masson-Pevet, M., and Hicks, D. (2006). Photoreceptor organization and rhythmic phagocytosis in the nile rat Arvicanthis ansorgei: a novel diurnal rodent model for the study of cone pathophysiology. Invest. Ophthalmol. Vis. Sci. 47, 3109–3118..

Bosch, E., Horwitz, J., and Bok, D. (1993). Phagocytosis of outer segments by retinal pigment epithelium: phagosome-lysosome interaction. J. Histochem. Cytochem. 41, 253–263..

Brenna, A., Olejniczak, I., Chavan, R., Ripperger, J.A., Langmesser, S., Cameroni, E., Hu, Z., De Virgilio, C., Dengjel, J., and Albrecht, U. (2019). Cyclin-dependent kinase 5 (CDK5) regulates the circadian clock. Elife 8. https://doi.org/10.7554/eLife.50925.

Chergui, K., Svenningsson, P., and Greengard, P. (2004). Cyclin-dependent kinase 5 regulates dopaminergic and glutamatergic transmission in the striatum. Proc. Natl. Acad. Sci. U. S. A. 101, 2191– 2196..

Contreras-Vallejos, E., Utreras, E., and Gonzalez-Billault, C. (2012). Going out of the brain: non-nervous system physiological and pathological functions of Cdk5. Cell. Signal. 24, 44–52..

Cottrell, G.T., Lin, R., Warn-Cramer, B.J., Lau, A.F., and Burt, J.M. (2003). Mechanism of v-Src- and mitogen-activated protein kinase-induced reduction of gap junction communication. Am. J. Physiol. Cell Physiol. 284, C511–C520..

Del Pozo, M.A., Kiosses, W.B., Alderson, N.B., Meller, N., Hahn, K.M., and Schwartz, M.A. (2002). Integrins regulate GTP-Rac localized effector interactions through dissociation of Rho-GDI. Nat. Cell Biol. 4, 232–239..

Dinet, V., Ansari, N., Torres-Farfan, C., and Korf, H.-W. (2007). Clock gene expression in the retina of melatonin-proficient (C3H) and melatonin-deficient (C57BL) mice. J. Pineal Res. 42, 83–91..

Dosch, M., Zindel, J., Jebbawi, F., Melin, N., Sanchez-Taltavull, D., Stroka, D., Candinas, D., and Beldi, G. (2019). Connexin-43-dependent ATP release mediates macrophage activation during sepsis. Elife 8. https://doi.org/10.7554/eLife.42670.

Dunn, C.A., and Lampe, P.D. (2014). Injury-triggered Akt phosphorylation of Cx43: a ZO-1-driven molecular switch that regulates gap junction size. J. Cell Sci. 127, 455–464..

Fadjukov, J., Wienbar, S., Hakanen, S., Aho, V., Vihinen-Ranta, M., Ihalainen, T.O., Schwartz, G.W., and Nymark, S. (2022). Gap junctions and connexin hemichannels both contribute to the electrical properties of retinal pigment epithelium. Journal of General Physiology 154. https://doi.org/10.1085/jgp.202112916.

Falk, M.M., Kells, R.M., and Berthoud, V.M. (2014). Degradation of connexins and gap junctions. FEBS Lett. 588, 1221–1229..

Finnemann, S.C., and Rodriguez-Boulan, E. (1999). Macrophage and retinal pigment epithelium phagocytosis: apoptotic cells and photoreceptors compete for alphavbeta3 and alphavbeta5 integrins, and protein kinase C regulates alphavbeta5 binding and cytoskeletal linkage. J. Exp. Med. 190, 861–874..

Fisher, S.K., Pfeffer, B.A., and Anderson, D.H. (1983). Both rod and cone disc shedding are related to light onset in the cat. Invest. Ophthalmol. Vis. Sci. 24, 844–856..

Flannery, J.G., and Fisher, S.K. (1984). Circadian disc shedding in Xenopus retina in vitro. Invest. Ophthalmol. Vis. Sci. 25, 229–232..

Fu, W.-Y., Chen, Y., Sahin, M., Zhao, X.-S., Shi, L., Bikoff, J.B., Lai, K.-O., Yung, W.-H., Fu, A.K.Y., Greenberg, M.E., et al. (2007). Cdk5 regulates EphA4-mediated dendritic spine retraction through an ephexin1-dependent mechanism. Nat. Neurosci. 10, 67–76..

Gabriel, R., Lesauter, J., Silver, R., Garcia-España, A., and Witkovsky, P. (2001). Diurnal and circadian variation of protein kinase C immunoreactivity in the rat retina. J. Comp. Neurol. 439, 140–150..

Glass, A.M., Wolf, B.J., Schneider, K.M., Princiotta, M.F., and Taffet, S.M. (2013). Connexin43 is dispensable for phagocytosis. J. Immunol. 190, 4830–4835..

Goel, M., and Mangel, S.C. (2021). Dopamine-Mediated Circadian and Light/Dark-Adaptive Modulation of Chemical and Electrical Synapses in the Outer Retina. Front. Cell. Neurosci. 15, 647541..

Goodenough, D.A., and Paul, D.L. (2009). Gap junctions. Cold Spring Harb. Perspect. Biol. 1, a002576..

Goodenough, D.A., Goliger, J.A., and Paul, D.L. (1996). Connexins, connexons, and intercellular communication. Annu. Rev. Biochem. 65, 475–502..

Goyal, V., DeVera, C., Laurent, V., Sellers, J., Chrenek, M.A., Hicks, D., Baba, K., Iuvone, P.M., and Tosini, G. (2020). Dopamine 2 Receptor Signaling Controls the Daily Burst in Phagocytic Activity in the Mouse Retinal Pigment Epithelium. Invest. Ophthalmol. Vis. Sci. 61, 10..

Hall, M.O., Abrams, T.A., and Mittag, T.W. (1993). The phagocytosis of rod outer segments is inhibited by drugs linked to cyclic adenosine monophosphate production. Invest. Ophthalmol. Vis. Sci. 34, 2392– 2401..

Hongisto, H., Jylhä, A., Nättinen, J., Rieck, J., Ilmarinen, T., Veréb, Z., Aapola, U., Beuerman, R., Petrovski, G., Uusitalo, H., et al. (2017). Comparative proteomic analysis of human embryonic stem cell-derived and primary human retinal pigment epithelium. Sci. Rep. 7, 6016..

Hossain, M.Z., Ao, P., and Boynton, A.L. (1998). Platelet-derived growth factor-induced disruption of gap junctional communication and phosphorylation of connexin43 involves protein kinase C and mitogen-activated protein kinase. J. Cell. Physiol. 176, 332–341..

Hunter, A.W., Barker, R.J., Zhu, C., and Gourdie, R.G. (2005). Zonula occludens-1 alters connexin43 gap junction size and organization by influencing channel accretion. Mol. Biol. Cell 16, 5686–5698..

Johansson, J.K., Karema-Jokinen, V.I., Hakanen, S., Jylhä, A., Uusitalo, H., Vihinen-Ranta, M., Skottman, H., Ihalainen, T.O., and Nymark, S. (2019). Sodium channels enable fast electrical signaling and regulate phagocytosis in the retinal pigment epithelium. BMC Biol. 17, 63..

Jones, J.C.R. (2016). Pre- and Post-embedding Immunogold Labeling of Tissue Sections. Methods Mol. Biol. 1474, 291–307..

Jordan, K., Chodock, R., Hand, A.R., and Laird, D.W. (2001). The origin of annular junctions: a mechanism of gap junction internalization. J. Cell Sci. 114, 763–773..

Kameritsch, P., Pogoda, K., and Pohl, U. (2012). Channel-independent influence of connexin 43 on cell migration. Biochim. Biophys. Acta 1818, 1993–2001..

Kameritsch, P., Kiemer, F., Beck, H., Pohl, U., and Pogoda, K. (2015). Cx43 increases serum induced filopodia formation via activation of p21-activated protein kinase 1. Biochim. Biophys. Acta 1853, 2907– 2917..

Ko, G.Y.-P., Shi, L., and Ko, M.L. (2009). Circadian regulation of ion channels and their functions. J. Neurochem. 110, 1150–1169..

Korkka, I., Viheriälä, T., Juuti-Uusitalo, K., Uusitalo-Järvinen, H., Skottman, H., Hyttinen, J., and Nymark, S. (2019). Functional Voltage-Gated Calcium Channels Are Present in Human Embryonic Stem Cell-Derived Retinal Pigment Epithelium. Stem Cells Transl. Med. 8, 179–193..

Korshunov, K.S., Blakemore, L.J., and Trombley, P.Q. (2017). Dopamine: A Modulator of Circadian Rhythms in the Central Nervous System. Front. Cell. Neurosci. 11, 91..

Krizaj, D., Gábriel, R., Owen, W.G., and Witkovsky, P. (1998). Dopamine D2 receptor-mediated modulation of rod-cone coupling in the Xenopus retina. J. Comp. Neurol. 398, 529–538..

Kwak, Y., Jeong, J., Lee, S., Park, Y.-U., Lee, S.-A., Han, D.-H., Kim, J.-H., Ohshima, T., Mikoshiba, K., Suh, Y.-H., et al. (2013). Cyclin-dependent kinase 5 (Cdk5) regulates the function of CLOCK protein by direct phosphorylation. J. Biol. Chem. 288, 36878–36889..

Laird, D.W., Puranam, K.L., and Revel, J.P. (1991). Turnover and phosphorylation dynamics of connexin43 gap junction protein in cultured cardiac myocytes. Biochem. J 273(Pt 1), 67–72..

Lampe, P.D. (1994). Analyzing phorbol ester effects on gap junctional communication: a dramatic inhibition of assembly. J. Cell Biol. 127, 1895–1905..

Lampe, P.D., and Lau, A.F. (2004). The effects of connexin phosphorylation on gap junctional communication. Int. J. Biochem. Cell Biol. 36, 1171–1186..

Lampe, P.D., TenBroek, E.M., Burt, J.M., Kurata, W.E., Johnson, R.G., and Lau, A.F. (2000). Phosphorylation of connexin43 on serine368 by protein kinase C regulates gap junctional communication. J. Cell Biol. 149, 1503–1512..

Lastwika, K.J., Dunn, C.A., Solan, J.L., and Lampe, P.D. (2019). Phosphorylation of connexin 43 at MAPK, PKC or CK1 sites each distinctly alter the kinetics of epidermal wound repair. J. Cell Sci. 132. https://doi.org/10.1242/jcs.234633.

LaVail, M.M. (1976). Rod outer segment disk shedding in rat retina: relationship to cyclic lighting. Science 194, 1071–1074..

LaVail, M.M. (1980). Circadian nature of rod outer segment disc shedding in the rat. Invest. Ophthalmol. Vis. Sci. 19, 407–411..

Leithe, E., Mesnil, M., and Aasen, T. (2018). The connexin 43 C-terminus: A tail of many tales. Biochim. Biophys. Acta Biomembr. 1860, 48–64..

Levin, M. (2007). Gap junctional communication in morphogenesis. Prog. Biophys. Mol. Biol. 94, 186– 206..

Li, H., Zhang, Z., Blackburn, M.R., Wang, S.W., Ribelayga, C.P., and O’Brien, J. (2013). Adenosine and Dopamine Receptors Coregulate Photoreceptor Coupling via Gap Junction Phosphorylation in Mouse Retina. Journal of Neuroscience 33, 3135–3150. https://doi.org/10.1523/jneurosci.2807-12.2013.

Liu, R., Tian, B., Gearing, M., Hunter, S., Ye, K., and Mao, Z. (2008). Cdk5-mediated regulation of the PIKE-A-Akt pathway and glioblastoma cell invasion. Proc. Natl. Acad. Sci. U. S. A. 105, 7570–7575..

Malfait, M., Gomez, P., van Veen, T.A., Parys, J.B., De Smedt, H., Vereecke, J., and Himpens, B. (2001). Effects of hyperglycemia and protein kinase C on connexin43 expression in cultured rat retinal pigment epithelial cells. J. Membr. Biol. 181, 31–40..

Mamaeva, D., Jazouli, Z., DiFrancesco, M.L., Erkilic, N., Dubois, G., Hilaire, C., Meunier, I., Boukhaddaoui, H., and Kalatzis, V. (2021). Novel roles for voltage-gated T-type Ca2+ and ClC-2 channels in phagocytosis and angiogenic factor balance identified in human iPSC-derived RPE. FASEB J. 35, e21406..

Mao, Y., and Finnemann, S.C. (2012). Essential diurnal Rac1 activation during retinal phagocytosis requires αvβ5 integrin but not tyrosine kinases focal adhesion kinase or Mer tyrosine kinase. Mol. Biol. Cell 23, 1104–1114..

Mao, Y., and Finnemann, S.C. (2013). Analysis of photoreceptor outer segment phagocytosis by RPE cells in culture. Methods Mol. Biol. 935, 285–295..

Mazzoni, F., Safa, H., and Finnemann, S.C. (2014). Understanding photoreceptor outer segment phagocytosis: use and utility of RPE cells in culture. Exp. Eye Res. 126, 51–60..

Mese, G., Richard, G., and White, T.W. (2007). Gap junctions: basic structure and function. J. Invest. Dermatol. 127, 2516–2524..

Milićević, N., Ait-Hmyed Hakkari, O., Bagchi, U., Sandu, C., Jongejan, A., Moerland, P.D., Ten Brink, J.B., Hicks, D., Bergen, A.A., and Felder-Schmittbuhl, M.-P. (2021). Core circadian clock genes Per1 and Per2 regulate the rhythm in photoreceptor outer segment phagocytosis. FASEB J. 35, e21722..

Müller, C., Más Gómez, N., Ruth, P., and Strauss, O. (2014). CaV1.3 L-type channels, maxiK Ca(2+)-dependent K(+) channels and bestrophin-1 regulate rhythmic photoreceptor outer segment phagocytosis by retinal pigment epithelial cells. Cell. Signal. 26, 968–978..

Musil, L.S., Beyer, E.C., and Goodenough, D.A. (1990). Expression of the gap junction protein connexin43 in embryonic chick lens: molecular cloning, ultrastructural localization, and post-translational phosphorylation. J. Membr. Biol. 116, 163–175..

Nakaya, M., Kitano, M., Matsuda, M., and Nagata, S. (2008). Spatiotemporal activation of Rac1 for engulfment of apoptotic cells. Proc. Natl. Acad. Sci. U. S. A. 105, 9198–9203..

Nandrot, E.F., Kim, Y., Brodie, S.E., Huang, X., Sheppard, D., and Finnemann, S.C. (2004). Loss of synchronized retinal phagocytosis and age-related blindness in mice lacking alphavbeta5 integrin. J. Exp. Med. 200, 1539–1545..

Nimlamool, W., Andrews, R.M.K., and Falk, M.M. (2015). Connexin43 phosphorylation by PKC and MAPK signals VEGF-mediated gap junction internalization. Mol. Biol. Cell 26, 2755–2768..

Pogoda, K., Kameritsch, P., Retamal, M.A., and Vega, J.L. (2016). Regulation of gap junction channels and hemichannels by phosphorylation and redox changes: a revision. BMC Cell Biol. 17 Suppl 1, 11..

Qi, G.-J., Chen, Q., Chen, L.-J., Shu, Y., Bu, L.-L., Shao, X.-Y., Zhang, P., Jiao, F.-J., Shi, J., and Tian, B. (2016). Phosphorylation of Connexin 43 by Cdk5 Modulates Neuronal Migration During Embryonic Brain Development. Mol. Neurobiol. 53, 2969–2982..

Rela, L., and Szczupak, L. (2004). Gap junctions: their importance for the dynamics of neural circuits. Mol. Neurobiol. 30, 341–357..

Rhett, J.M., Jourdan, J., and Gourdie, R.G. (2011). Connexin 43 connexon to gap junction transition is regulated by zonula occludens-1. Mol. Biol. Cell 22, 1516–1528..

Ribelayga, C., Cao, Y., and Mangel, S.C. (2008). The circadian clock in the retina controls rod-cone coupling. Neuron 59, 790–801..

Schneider, C.A., Rasband, W.S., and Eliceiri, K.W. (2012). NIH Image to ImageJ: 25 years of image analysis. Nat. Methods 9, 671–675..

Segretain, D., and Falk, M.M. (2004). Regulation of connexin biosynthesis, assembly, gap junction formation, and removal. Biochim. Biophys. Acta 1662, 3–21..

Shah, K., and Rossie, S. (2018). Tale of the Good and the Bad Cdk5: Remodeling of the Actin Cytoskeleton in the Brain. Mol. Neurobiol. 55, 3426–3438..

Skottman, H. (2010). Derivation and characterization of three new human embryonic stem cell lines in Finland. In Vitro Cell. Dev. Biol. Anim. 46, 206–209..

Solan, J.L., and Lampe, P.D. (2008). Connexin 43 in LA-25 cells with active v-src is phosphorylated on Y247, Y265, S262, S279/282, and S368 via multiple signaling pathways. Cell Commun. Adhes. 15, 75– 84..

Solan, J.L., and Lampe, P.D. (2014). Specific Cx43 phosphorylation events regulate gap junction turnover in vivo. FEBS Lett. 588, 1423–1429..

Solan, J.L., and Lampe, P.D. (2018). Spatio-temporal regulation of connexin43 phosphorylation and gap junction dynamics. Biochim. Biophys. Acta Biomembr. 1860, 83–90..

Sorgen, P.L., Duffy, H.S., Sahoo, P., Coombs, W., Delmar, M., and Spray, D.C. (2004). Structural changes in the carboxyl terminus of the gap junction protein connexin43 indicates signaling between binding domains for c-Src and zonula occludens-1. J. Biol. Chem. 279, 54695–54701..

Sorgen, P.L., Trease, A.J., Spagnol, G., Delmar, M., and Nielsen, M.S. (2018). Protein^-^Protein Interactions with Connexin 43: Regulation and Function. Int. J. Mol. Sci. 19. https://doi.org/10.3390/ijms19051428.

Strauss, O. (2005). The retinal pigment epithelium in visual function. Physiol. Rev. 85, 845–881..

Sutor, B., and Hagerty, T. (2005). Involvement of gap junctions in the development of the neocortex. Biochim. Biophys. Acta 1719, 59–68..

Tan, T.C., Valova, V.A., Malladi, C.S., Graham, M.E., Berven, L.A., Jupp, O.J., Hansra, G., McClure, S.J., Sarcevic, B., Boadle, R.A., et al. (2003). Cdk5 is essential for synaptic vesicle endocytosis. Nat. Cell Biol. 5, 701–710..

de Thonel, A., Ferraris, S.E., Pallari, H.-M., Imanishi, S.Y., Kochin, V., Hosokawa, T., Hisanaga, S.-I., Sahlgren, C., and Eriksson, J.E. (2010). Protein kinase Czeta regulates Cdk5/p25 signaling during myogenesis. Mol. Biol. Cell 21, 1423–1434..

Tomizawa, K., Sunada, S., Lu, Y.-F., Oda, Y., Kinuta, M., Ohshima, T., Saito, T., Wei, F.-Y., Matsushita, M., Li, S.-T., et al. (2003). Cophosphorylation of amphiphysin I and dynamin I by Cdk5 regulates clathrin-mediated endocytosis of synaptic vesicles. J. Cell Biol. 163, 813–824..

Toyofuku, T., Akamatsu, Y., Zhang, H., Kuzuya, T., Tada, M., and Hori, M. (2001). c-Src regulates the interaction between connexin-43 and ZO-1 in cardiac myocytes. J. Biol. Chem. 276, 1780–1788..

Trenholm, S., and Awatramani, G.B. (2019). Myriad roles for gap junctions in retinal circuits. In Webvision: The Organization of the Retina and Visual System, H. Kolb, E. Fernandez, and R. Nelson, eds. (Salt Lake City (UT): University of Utah Health Sciences Center),.

Uribe-Querol, E., and Rosales, C. (2020). Phagocytosis: Our Current Understanding of a Universal Biological Process. Front. Immunol. 11, 1066..

Vaajasaari, H., Ilmarinen, T., Juuti-Uusitalo, K., Rajala, K., Onnela, N., Narkilahti, S., Suuronen, R., Hyttinen, J., Uusitalo, H., and Skottman, H. (2011). Toward the defined and xeno-free differentiation of functional human pluripotent stem cell-derived retinal pigment epithelial cells. Mol. Vis. 17, 558–575..

Vargas, J.A., and Finnemann, S.C. (2022). Differences in Diurnal Rhythm of Rod Outer Segment Renewal between 129T2/SvEmsJ and C57BL/6J Mice. Int. J. Mol. Sci. 23. https://doi.org/10.3390/ijms23169466.

Veeraraghavan, R., Gourdie, R.G., and Poelzing, S. (2014). Mechanisms of cardiac conduction: a history of revisions. Am. J. Physiol. Heart Circ. Physiol. 306, H619–H627..

Viheriälä, T., Sorvari, J., Ihalainen, T.O., Mörö, A., Grönroos, P., Schlie-Wolter, S., Chichkov, B., Skottman, H., Nymark, S., and Ilmarinen, T. (2021). Culture surface protein coatings affect the barrier properties and calcium signalling of hESC-RPE. Sci. Rep. 11, 1–14..

Völgyi, B., Kovács-Oller, T., Atlasz, T., Wilhelm, M., and Gábriel, R. (2013). Gap junctional coupling in the vertebrate retina: variations on one theme? Prog. Retin. Eye Res. 34, 1–18..

Wavre-Shapton, S.T., Meschede, I.P., Seabra, M.C., and Futter, C.E. (2014). Phagosome maturation during endosome interaction revealed by partial rhodopsin processing in retinal pigment epithelium. J. Cell Sci. 127, 3852–3861..

White, T.W., and Paul, D.L. (1999). Genetic diseases and gene knockouts reveal diverse connexin functions. Annu. Rev. Physiol. 61, 283–310..

Willecke, K., Eiberger, J., Degen, J., Eckardt, D., Romualdi, A., Güldenagel, M., Deutsch, U., and Söhl, G. (2002). Structural and functional diversity of connexin genes in the mouse and human genome. Biol. Chem. 383, 725–737..

Yang, M., James, A.D., Suman, R., Kasprowicz, R., Nelson, M., O’Toole, P.J., and Brackenbury, W.J. (2020). Voltage-dependent activation of Rac1 by Nav 1.5 channels promotes cell migration. J. Cell. Physiol. 235, 3950–3972..

Young, R.W. (1978). The daily rhythm of shedding and degradation of rod and cone outer segment membranes in the chick retina. Invest. Ophthalmol. Vis. Sci. 17, 105–116..

